# Calcium-independent biosensors for multiplexed in vivo imaging of extracellular lactate dynamics

**DOI:** 10.64898/2026.10.02.756141

**Authors:** Jia-Jin Liu, Hidenobu Mizuno, Emmanuel Than-Trong, Philipp Mächler, Hideaki Kubotera, Tony Barbay, Ching-Yao Wang, Qin-Ling To, Yuki Kamijo, Taki Nishimura, Marc Boisvert, Marie-Eve Paquet, Mikhail Drobizhev, Stephanie J. Cragg, Jun Nagai, David Kleinfeld, Serge Charpak, Yusuke Nasu

**Author notes:** These authors contributed equally to this work (Hidenobu Mizuno, Emmanuel Than-Trong, Philipp Mächler, Hideaki Kubotera, Tony Barbay). Correspondence should be addressed to: Yusuke Nasu.

## Abstract

Extracellular L-lactate reports cellular metabolism, but resolving its dynamics alongside intracellular metabolism and neuronal activity remains challenging. Here we develop eLACCO3 and R-eLACCO3, green and red cell-surface lactate biosensors with Ca^2+^-independent responses and faster response and recovery kinetics in cultured cells. Selected variants provide larger fluorescence responses than their predecessors in brain cells, slices, and living mice and improve detection of glucose-evoked extracellular lactate oscillations. Combining R-eLACCO3 with GCaMP6f enabled imaging of extracellular L-lactate simultaneously with odor-evoked activity in anesthetized mice and cortical activity in awake mice. These biosensors enable multiplexed measurements of extracellular lactate dynamics in relation to intracellular metabolism and neuronal activity.

## Introduction

Intercellular L-lactate exchange contributes to metabolic cooperation and signaling in physiology and disease^1–5^. In the brain, extracellular lactate measurements complement intracellular readouts by reporting lactate availability between cells. Comparing these measurements with neuronal activity could help establish how extracellular lactate dynamics relate to changing neural demand across behavioral transitions.

Genetically encoded fluorescent biosensors have enabled intracellular lactate imaging^6–15^, and green eLACCO^14,16^ and red R-eLACCO^17^ biosensors have extended these measurements to the extracellular space. However, the relatively slow response and recovery kinetics and limited fluorescence responses of existing extracellular biosensors can obscure transient or low-amplitude signals. Their Ca^2+^-dependent lactate recognition introduces an additional potential confound because extracellular Ca^2+^ changes with neuronal activity^18^. These limitations are particularly consequential when comparing extracellular lactate signals with intracellular metabolism or neuronal Ca^2+^ dynamics. Ca^2+^-independent extracellular biosensors with improved temporal responses and signal changes are therefore needed, together with complementary fluorescence colors for multiplexed imaging.

Here we engineered eLACCO3 and R-eLACCO3, green and red extracellular lactate biosensors, through rational mutagenesis, directed evolution, and optimization of cell-surface targeting. We characterized their Ca^2+^ independence and response kinetics in cultured cells and compared their performance with that of previous extracellular sensors in cultured cells, brain slices, and living mice. The resulting biosensors improved detection of endogenous extracellular lactate oscillations in cultured cells. In the mouse brain, pairing R-eLACCO3 with GCaMP6f enabled simultaneous measurements of extracellular lactate and neuronal Ca^2+^ signals.

## Results

### Development and characterization of Ca^2+^-independent extracellular lactate biosensors

To develop Ca^2+^-independent extracellular L-lactate biosensors, we selected the LldR-based iLACCO1 and R-iLACCO1 as starting scaffolds rather than Ca^2+^-dependent TTHA0766-derived sensors (**Fig. 1a**) ^14,15,17^. To minimize disulfide formation under oxidizing conditions^19^, we introduced Cys53Ala in LldR, together with Cys257Ser in cpGFP or Cys180Tyr in cpmApple. The resulting iLACCO1.3 and R-iLACCO1.3 retained lactate responses comparable to those of their parental sensors (**Supplementary Fig. 1a**). We then performed iterative directed evolution using lysate-based screening for lactate-induced Δ*F*/*F* (Fig. 1a and **Supplementary Fig. 1b**). Six rounds yielded iLACCO1.4 (lysate Δ*F*/*F* = 20). Insertion of a Trp110–Ser111 linker between LldR and cpGFP initially reduced responses, but further evolution produced iLACCO2_H_ (lysate Δ*F*/*F* = 36; 30 mutations relative to iLACCO1; **Supplementary Fig. 2**). In parallel, seven rounds yielded R-iLACCO1.9 (lysate Δ*F*/*F* = 34), and combining Ala141Val, Gln154Pro, and Arg300Ser produced R-iLACCO2_H_, with 13 substitutions relative to R-iLACCO1 (**Supplementary Fig. 3**). Additional mutagenesis generated medium- and low-affinity variants and lactate-insensitive controls (Supplementary Figs. 2,3). Detailed characterization of the iLACCO2 and R-iLACCO2 variants as purified proteins and in live mammalian cells is provided in **Supplementary Figs. 4–6**, **Supplementary Tables 1,2**, and **Supplementary Note 1**.

**Fig. 1.**
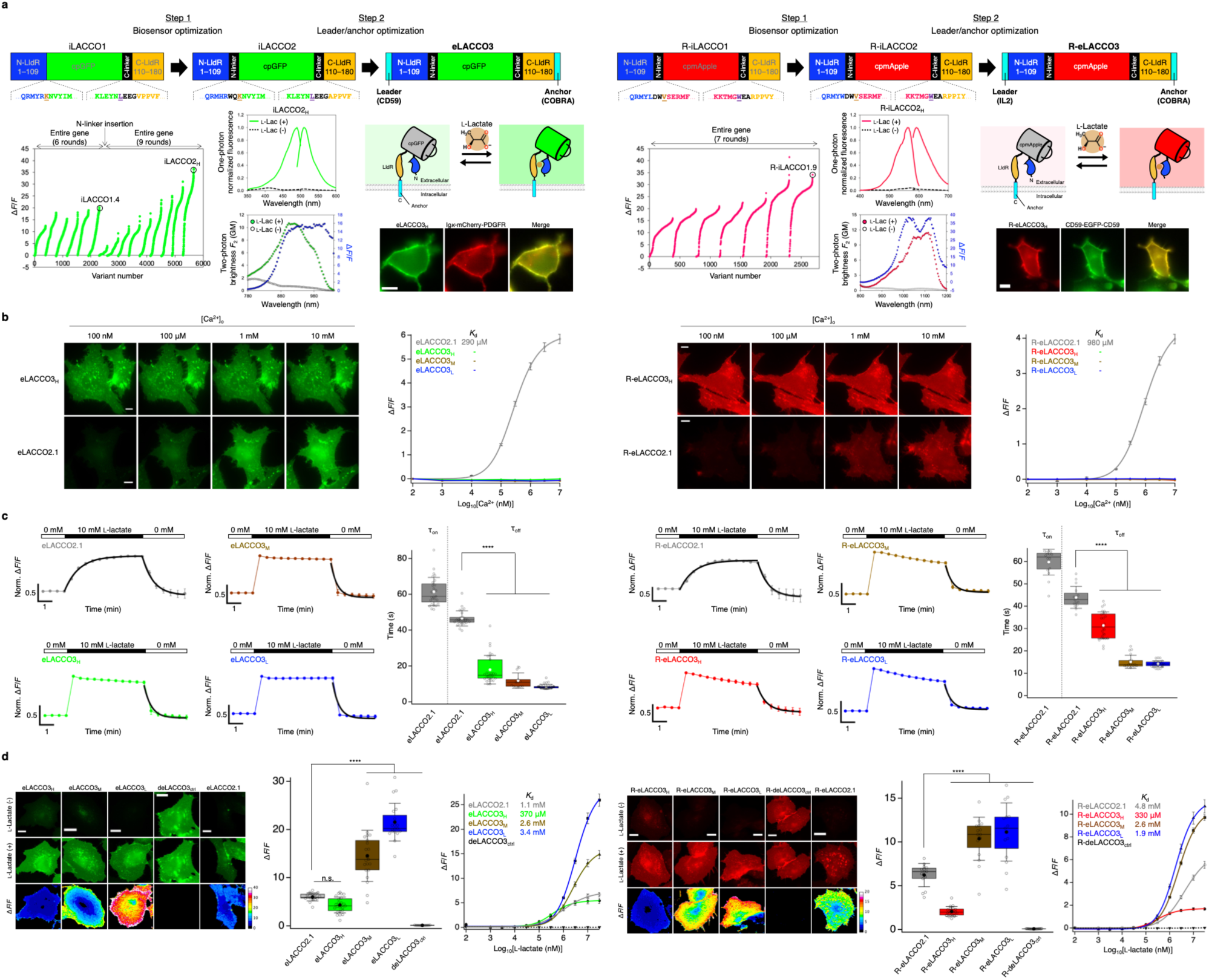
Development and characterization of Ca^2+^-independent green and red biosensors for extracellular lactate. (**a**) The strategy for developing eLACCO3 and R-eLACCO3 biosensors. In the primary structures, linker regions are shown in black and the two “gate post” residues^36^ in cpFP are highlighted in dark orange and purple. Δ*F*/*F* rank plot representing all proteins tested during the directed evolution. Graphs indicate one-photon (top) and two-photon (bottom) spectra of purified iLACCO2_H_ and R-iLACCO2_H_ in the presence (10 mM) and absence of L-lactate. Cell images show the localization of eLACCO3_H_ and R-eLACCO3_H_ in HEK293T cells. Scale bars, 10 μm. (**b**) Ca^2+^ titration of eLACCO and R-eLACCO variants on HeLa cells in the presence (10 mM) of L-lactate. Mean ± s.e.m. *n* = 39, 39, 23, 42, 39, 50, 41, and 46 cells for eLACCO2.1, eLACCO3_H_, eLACCO3_M_, eLACCO3_L_, R-eLACCO2.1, R-eLACCO3_H_, R-eLACCO3_M_, and R-eLACCO3_L_, respectively. Scale bars, 10 μm. (**c**) Time course (left, mean ± s.d.) and time constant (right) of the fluorescence response of eLACCO and R-eLACCO variants on HeLa cells upon L-lactate (10 mM) treatment. *n* = 27, 29, 13, 36, 16, 26, 20, and 26 cells for eLACCO2.1, eLACCO3_H_, eLACCO3_M_, eLACCO3_L_, R-eLACCO2.1, R-eLACCO3_H_, R-eLACCO3_M_, and R-eLACCO3_L_, respectively. One-way ANOVA with the Dunnett’s post hoc tests. \*\*\*\**p* < 0.0001. (**d**) Fluorescence images (left), Δ*F*/*F* for 10 mM L-lactate (middle), and L-lactate titration (right, mean ± s.e.m.) of eLACCO and R-eLACCO variants on HeLa cells. Scale bars, 10 μm. *n* = 18, 23, 21, 19, 13, 14, 19, 16, 15, and 19 cells for eLACCO2.1, eLACCO3_H_, eLACCO3_M_, eLACCO3_L_, deLACCO3_ctrl_, R-eLACCO2.1, R-eLACCO3_H_, R-eLACCO3M, R-eLACCO3_L_, and R-deLACCO3_ctrl_, respectively. One-way ANOVA with the Dunnett’s post hoc tests. *p* (eLACCO2.1 vs. eLACCO3_H_) = 0.2854, \*\*\*\**p* < 0.0001.

To enable extracellular L-lactate sensing, we optimized cell-surface targeting using N-terminal leader sequences and C-terminal membrane anchors. Because targeting sequences affect sensor performance^14,17^, we screened anchors using iLACCO1.1 (ref. ^15^) and R-iLACCO1.2 (ref. ^14^) fused to a human CD59 leader. We assessed surface localization in HEK293T cells and basal fluorescence and Δ*F*/*F* in HeLa cells (**Supplementary Fig. 7a,b**). The PDGFR anchor and COBRA-derived glycosylphosphatidylinositol (GPI)-anchoring sequence were selected for screening with 11 leader sequences. COBRA generally supported better surface localization than PDGFR. The CD59 leader provided a balance of localization, basal fluorescence, and response for iLACCO1.1, whereas the interleukin-2 (IL2) leader produced the largest R-iLACCO1.2 responses, although localization and basal fluorescence were lower (**Supplementary Fig. 7c**). Applying these targeting configurations to the engineered sensor cores generated CD59–iLACCO2–COBRA and IL2–R-iLACCO2–COBRA constructs, designated eLACCO3 and R-eLACCO3, respectively (Fig. 1a).

All variants localized to the plasma membrane (Fig. 1a and **Supplementary Fig. 7d**). Both series showed Ca^2+^-independent fluorescence over the tested Ca^2+^ range in the presence of 10 mM L-lactate, unlike their TTHA0766-based predecessors (**Fig. 1b**)^14,16,17^. Following lactate addition, all eLACCO3 and R-eLACCO3 variants approached maximal fluorescence by the first measurement at 30 s, precluding reliable estimation of their response time constants (τ_on_). By contrast, eLACCO2.1 and R-eLACCO2.1 exhibited τ_on_ values of 61.5 ± 0.3 s and 59.7 ± 1.4 s, respectively (mean ± s.e.m.). Recovery after lactate removal was also faster for both engineered series. Apparent recovery time constants (τ_off_) ranged from 8.5 to 17.9 s for eLACCO3 variants and from 14.2 to 31.4 s for R-eLACCO3 variants, compared with 46.5 ± 0.2 s for eLACCO2.1 and 44.0 ± 1.2 s for R-eLACCO2.1 (**Fig. 1c**).

In HeLa cells, 10 mM L-lactate elicited Δ*F*/*F* values of 4.3 ± 0.3, 14.5 ± 1.2, and 21.5 ± 0.9 for eLACCO3_H_, eLACCO3_M_, and eLACCO3_L_, respectively, compared with 6.0 ± 0.2 for eLACCO2.1. Corresponding responses for R-eLACCO3_H_, R-eLACCO3_M_, and R-eLACCO3_L_ were 2.1 ± 0.1, 10.4 ± 0.6, and 11.1 ± 0.9, compared with 6.2 ± 0.4 for R-eLACCO2.1. The M and L variants therefore produced 2.4- and 3.6-fold larger responses than eLACCO2.1 in the green series, and 1.7- and 1.8-fold larger responses than R-eLACCO2.1 in the red series. The surface-localized controls deLACCO3_ctrl_ and R-deLACCO3_ctrl_ showed minimal responses (**Fig. 1d**). Apparent in situ *K*_d_ values were 370 μM, 2.6 mM, and 3.4 mM for eLACCO3_H_, eLACCO3_M_, and eLACCO3_L_, respectively, and 330 μM, 2.6 mM, and 1.9 mM for R-eLACCO3_H_, R-eLACCO3_M_ and R-eLACCO3L, respectively (Fig. 1d).

Flow cytometry showed that basal cellular fluorescence was generally comparable to or higher than that of the respective predecessors (**Supplementary Fig. 7e**). Photostability varied among variants and with lactate availability. eLACCO3_M_ was more photostable than eLACCO2.1 in the absence of lactate, whereas eLACCO3_H_ and eLACCO3_L_ were less photostable both with and without lactate. R-eLACCO3_M_ and R-eLACCO3_L_ were more photostable than R-eLACCO3_H_, although R-eLACCO2.1 remained more photostable than all three engineered red variants (**Supplementary Fig. 7f**). Blue-light illumination increased red fluorescence, with larger changes in R-eLACCO3_H_ and R-deLACCO3_ctrl_ (**Supplementary Fig. 7g**).

### Improved detection of endogenous extracellular lactate dynamics

We next asked whether the improved sensor responses enabled better detection of endogenous extracellular lactate dynamics. To this end, we stimulated glucose-starved HeLa cells with glucose, which promotes lactate production and release (**Fig. 2a**)^14,17^. Fluorescence increased across both series. High-affinity variants responded most strongly: eLACCO3_H_ exhibited Δ*F*/*F* = 0.91 ± 0.05 versus 0.37 ± 0.03 for eLACCO2.1 (2.5-fold improvement), and R-eLACCO3_H_ exhibited 0.59 ± 0.05 versus 0.04 ± 0.01 for R-eLACCO2.1 (∼15-fold improvement). deLACCO3_ctrl_ responded minimally, whereas R-deLACCO3_ctrl_ fluorescence gradually declined, likely through photobleaching (**Fig. 2b**).

**Fig. 2.**
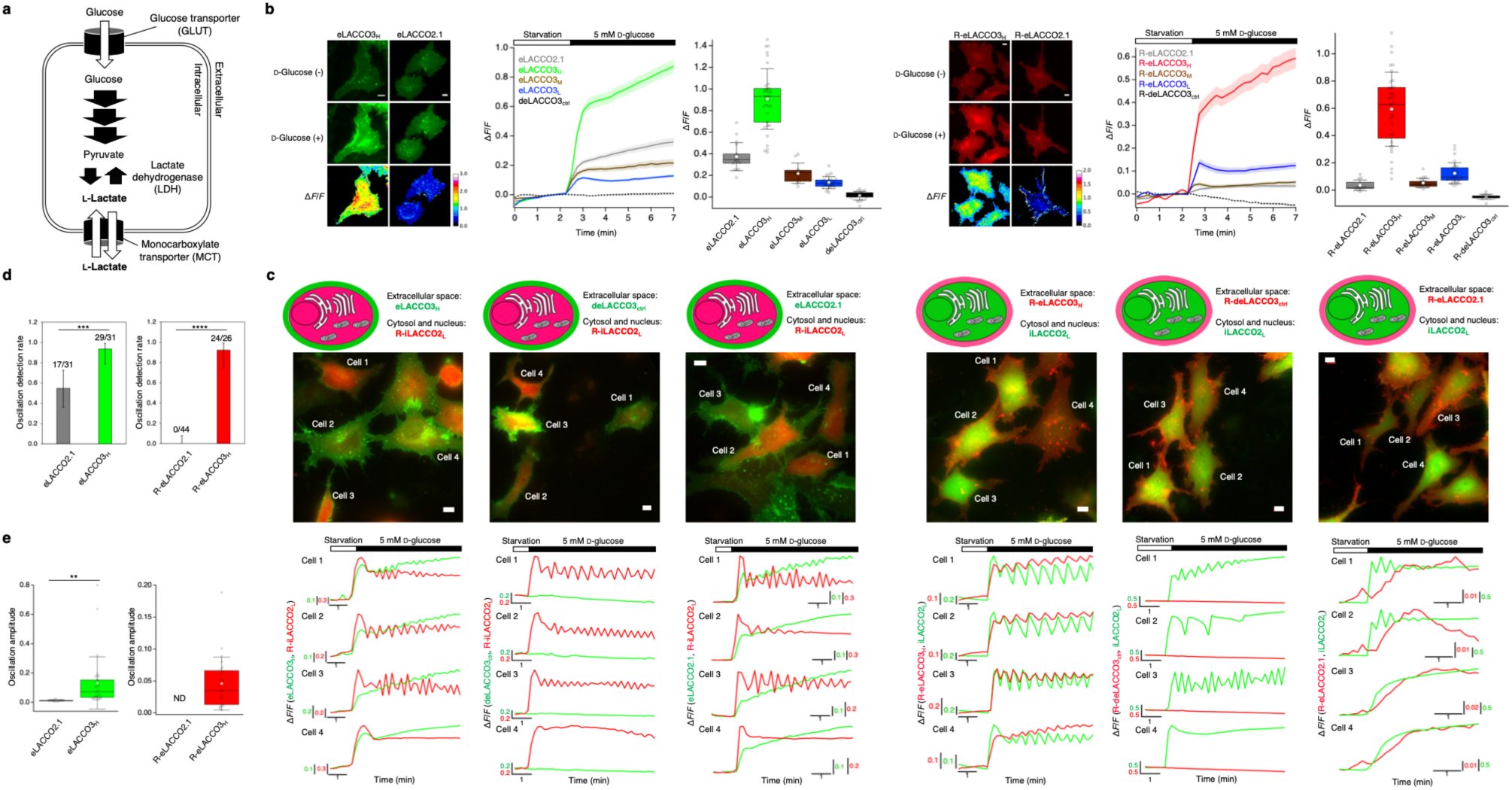
Improved detection of endogenous extracellular lactate dynamics. (**a**) Schematic representation of L-lactate metabolism upon D-glucose treatment. (**b**) Fluorescence images (left), traces (middle, mean ± s.e.m.), and Δ*F*/*F* (right) in HeLa cells upon 5 mM D-glucose treatment after starvation. Scale bars, 10 μm. *n* = 17, 34, 12, 25, 21, 12, 35, 21, 28, and 14 cells for eLACCO2.1, eLACCO3_H_, eLACCO3M, eLACCO3_L_, deLACCO3_ctrl_, R-eLACCO2.1, R-eLACCO3_H_, R-eLACCO3_M_, R-eLACCO3_L_, and R-deLACCO3_ctrl_, respectively. (**c**) Fluorescence imaging of lactate oscillation in HeLa cells upon 5 mM D-glucose treatment after starvation. Scale bars, 10 μm. (**d**) Detection rates of extracellular lactate oscillations in (c). Error bars indicate exact binomial 95% confidence intervals, and labels indicate detected cells / total cells that show intracellular lactate oscillation. Two-sided Fisher’s exact test. \*\*\**p* = 0.001, \*\*\*\**p* < 0.0001. (**e**) Oscillation amplitude of eLACCO and R-eLACCO variants. *n* = 17, 29, and 24 cells for eLACCO2.1, eLACCO3_H_, and R-eLACCO3_H_, respectively. Two-tailed student’s *t*-test with Welch’s correction, \*\**p* = 0.0013. ND, not detected.

To compare extracellular signals with intracellular lactate dynamics in individual cells, we paired eLACCO3_H_ with R-iLACCO2_L_ or R-eLACCO3_H_ with iLACCO2_L_ (**Fig. 2c**). Among cells exhibiting intracellular lactate oscillations^15^, eLACCO3_H_ detected extracellular oscillations in 29 of 31 cells, compared with 17 of 31 cells for eLACCO2.1. R-eLACCO3_H_ detected extracellular oscillations in 24 of 26 cells, whereas R-eLACCO2.1 detected none in 44 cells (**Fig. 2d**). Lactate-insensitive extracellular controls showed no oscillatory responses. Among cells with detectable extracellular oscillations, eLACCO3_H_ exhibited a larger mean fluorescence oscillation amplitude than eLACCO2.1 (Δ*F*/*F* = 0.13 ± 0.03 versus 0.01 ± 0.00). The corresponding mean amplitude for R-eLACCO3_H_ was 0.05 ± 0.01 (**Fig. 2e**).

Neighboring cells without intracellular oscillations also lacked detectable extracellular oscillations, consistent with spatially heterogeneous pericellular signals rather than a uniform change throughout the medium. Together, these comparisons demonstrate improved detection of endogenous extracellular lactate transients under the tested conditions.

### Validation of eLACCO3 and R-eLACCO3 in brain cells, brain slices, and living mice

Although eLACCO3_L_ and R-eLACCO3_L_ produced the largest HeLa Δ*F*/*F* upon exogenous L-lactate, their low basal brightness could compromise in vivo signal-to-noise ratios. Subsequent experiments therefore focused on the brighter high- and medium-affinity variants. In rat primary cortical neurons, human synapsin (hSyn)-driven sensors localized to membranes (**Fig. 3a**). At 10 mM L-lactate, eLACCO3_H_ and eLACCO3_M_ exhibited Δ*F*/*F* values of 22.2 ± 0.9 and 16.9 ± 2.0 versus 6.0 ± 0.4 for eLACCO2.1 (3.7- and 2.8-fold improvements); R-eLACCO3_H_ and R-eLACCO3_M_ exhibited 10.6 ± 1.3 and 10.4 ± 0.5 versus 4.8 ± 0.5 for R-eLACCO2.1 (∼2.2-fold improvements). In both cases, control constructs (deLACCO3_ctrl_ and R-deLACCO3_ctrl_) displayed no detectable response. In rat primary cortical astrocytes, gfaABC1D-driven sensors localized to the surface (**Fig. 3b**). eLACCO3_H_ and eLACCO3_M_ responses were 26.4 ± 1.4 and 22.4 ± 1.1 versus 7.5 ± 0.3 for eLACCO2.1; R-eLACCO3_H_ and R-eLACCO3_M_ responses were 11.5 ± 0.6 and 13.6 ± 1.3 versus 4.7 ± 0.6 for R-eLACCO2.1. Controls were unresponsive in both cell types.

**Fig. 3.**
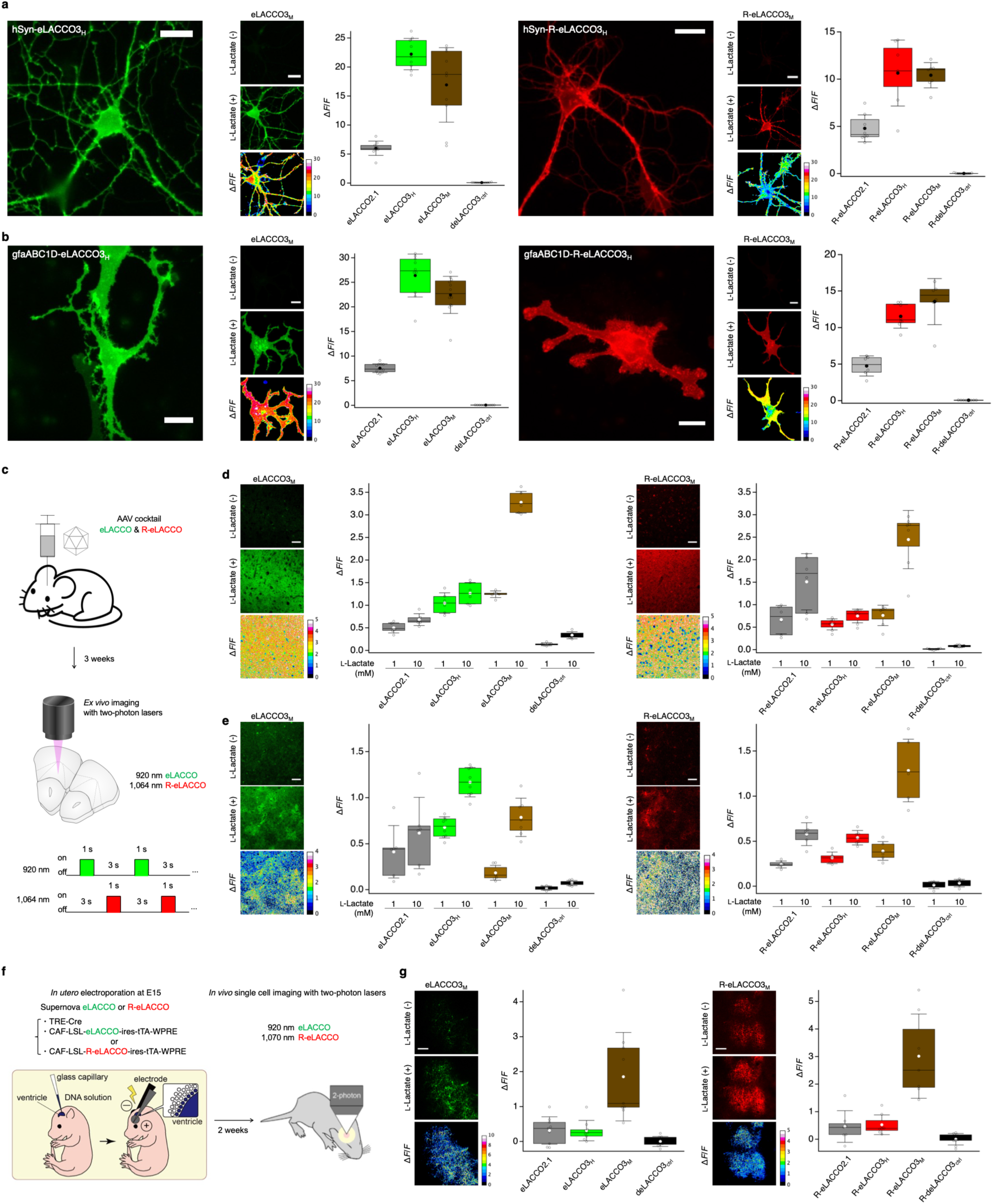
Validation of eLACCO3 and R-eLACCO3 in brain cells, brain slices, and living mice. (**a**) Fluorescence images (left) of primary cultured rat cortical neurons expressing eLACCO3_H_ or R-eLACCO3_H_. Fluorescence images (middle) and Δ*F*/*F* (right) of eLACCO and R-eLACCO variants expressed on cell surface of rat cortical neurons before and after 10 mM L-lactate stimulation. Scale bars, 20 μm. *n* = 9, 9, 10, 11, 8, 7, 7, and 10 cells for eLACCO2.1, eLACCO3_H_, eLACCO3_M_, deLACCO3_ctrl_, R-eLACCO2.1, R-eLACCO3_H_, R-eLACCO3_M_, and R-deLACCO3_ctrl_, respectively. (**b**) Fluorescence images (left) of primary cultured rat cortical astrocytes expressing eLACCO3_H_ and R-eLACCO3_H_. Fluorescence images (middle) and Δ*F*/*F* (right) of eLACCO and R-eLACCO variants expressed on cell surface of rat cortical astrocytes before and after 10 mM L-lactate stimulation. Scale bars, 20 μm. *n* = 12, 10, 12, 10, 6, 8, 6, and 8 cells for eLACCO2.1, eLACCO3_H_, eLACCO3_M_, deLACCO3_ctrl_, R-eLACCO2.1, R-eLACCO3_H_, R-eLACCO3_M_, and R-deLACCO3_ctrl_, respectively. (**c**) Schematic illustration of double AAV injection into the orbitofrontal cortex and fluorescence recording for the brain slices. (**d**) Fluorescence images (left) and L-lactate-dependent Δ*F*/*F* (right) of the brain slices expressing eLACCO and R-eLACCO variants under neuronal promoter hSyn before and after 10 mM L-lactate stimulation. Scale bars, 50 μm. *n* = 6, 6, 6, 6, 6, 8, 7, and 7 slices from 2, 2, 2, 2, 3, 3, 2, and 2 mice for eLACCO2.1, eLACCO3_H_, eLACCO3_M_, deLACCO3_ctrl_, R-eLACCO2.1, R-eLACCO3_H_, R-eLACCO3_M_, and R-deLACCO3_ctrl_, respectively. (**e**) Fluorescence images (left) and L-lactate-dependent Δ*F*/*F* (right) of the brain slices expressing eLACCO and R-eLACCO variants under astrocyte promoter gfaABC1D before and after 10 mM L-lactate stimulation. Scale bars, 20 μm. *n* = 8, 8, 7, 7, 6, 6, 6, and 6 slices from 3, 3, 2, 2, 2, 2, 2, and 2 mice for eLACCO2.1, eLACCO3_H_, eLACCO3_M_, deLACCO3_ctrl_, R-eLACCO2.1, R-eLACCO3_H_, R-eLACCO3_M_, and R-deLACCO3_ctrl_, respectively. (**f**) Schematic illustration of in utero electroporation into the ventricle of mouse embryos and fluorescence recording for the brain of living mouse pups. (**g**) Fluorescence images (left) and Δ*F*/*F* (right) of the brain in living mice expressing eLACCO and R-eLACCO variants before and after 10 mM L-lactate stimulation. Scale bars, 100 μm. *n* = 9, 9, 9, 9, 7, 8, 7, and 9 cells from 3 mice for eLACCO2.1, eLACCO3_H_, eLACCO3_M_, deLACCO3_ctrl_, R-eLACCO2.1, R-eLACCO3H, R-eLACCO3_M_, and R-deLACCO3_ctrl_, respectively.

We next expressed biosensors in the mouse orbitofrontal cortex using adeno-associated viruses (AAVs) under hSyn or gfaABC1D promoters and examined brain slices (**Fig. 3c**). Neuronal eLACCO3_H_ and eLACCO3_M_ responses exceeded those of eLACCO2.1 at both 1 and 10 mM L-lactate: Δ*F*/*F* values were 1.05 ± 0.09 and 1.24 ± 0.03 at 1 mM, and 1.27 ± 0.10 and 3.28 ± 0.10 at 10 mM, versus 0.50 ± 0.05 and 0.68 ± 0.05, respectively. Neuronal R-eLACCO3_H_ and R-eLACCO3_M_ responses at 1 mM (0.56 ± 0.04 and 0.76 ± 0.09) were comparable to R-eLACCO2.1 (0.67 ± 0.13). At 10 mM, the R-eLACCO3_M_ response reached 2.45 ± 0.24, exceeding R-eLACCO2.1 (1.51 ± 0.26), whereas the R-eLACCO3H response remained lower (0.75 ± 0.05; **Fig. 3d**). Astrocytic eLACCO3_H_ and eLACCO3_M_ responses were 0.68 ± 0.04 and 0.18 ± 0.03 at 1 mM, versus 0.41 ± 0.12 for eLACCO2.1; at 10 mM, both exceeded eLACCO2.1 (1.17 ± 0.06 and 0.79 ± 0.08 versus 0.62 ± 0.16). Astrocytic R-eLACCO3H and R-eLACCO3_M_ responses slightly exceeded R-eLACCO2.1 at 1 mM (0.32 ± 0.03 and 0.39 ± 0.04 versus 0.24 ± 0.02). At 10 mM, the R-eLACCO3_M_ response reached 1.28 ± 0.14, whereas R-eLACCO3H and R-eLACCO2.1 showed limited further increases (0.54 ± 0.03 and 0.58 ± 0.05; **Fig. 3e**). Controls showed negligible responses.

In living mice, biosensors were sparsely expressed in cortical layer 2/3 pyramidal neurons by in utero electroporation (**Fig. 3f**)^20^. Bath application of 10 mM L-lactate elicited Δ*F*/*F* values of 1.86 ± 0.42 and 3.01 ± 0.58 for eLACCO3_M_ and R-eLACCO3_M_, respectively, versus 0.32 ± 0.13 and 0.46 ± 0.22 for their predecessors (5.9- and 6.5-fold improvements). eLACCO3H and R-eLACCO3_H_ responses (0.31 ± 0.10 and 0.53 ± 0.13) were comparable to their respective predecessors; controls were unresponsive (**Fig. 3g**). **Table 1** summarizes the performance characteristics of eLACCO3 and R-eLACCO3 variants and their predecessors.

**Table 1.**
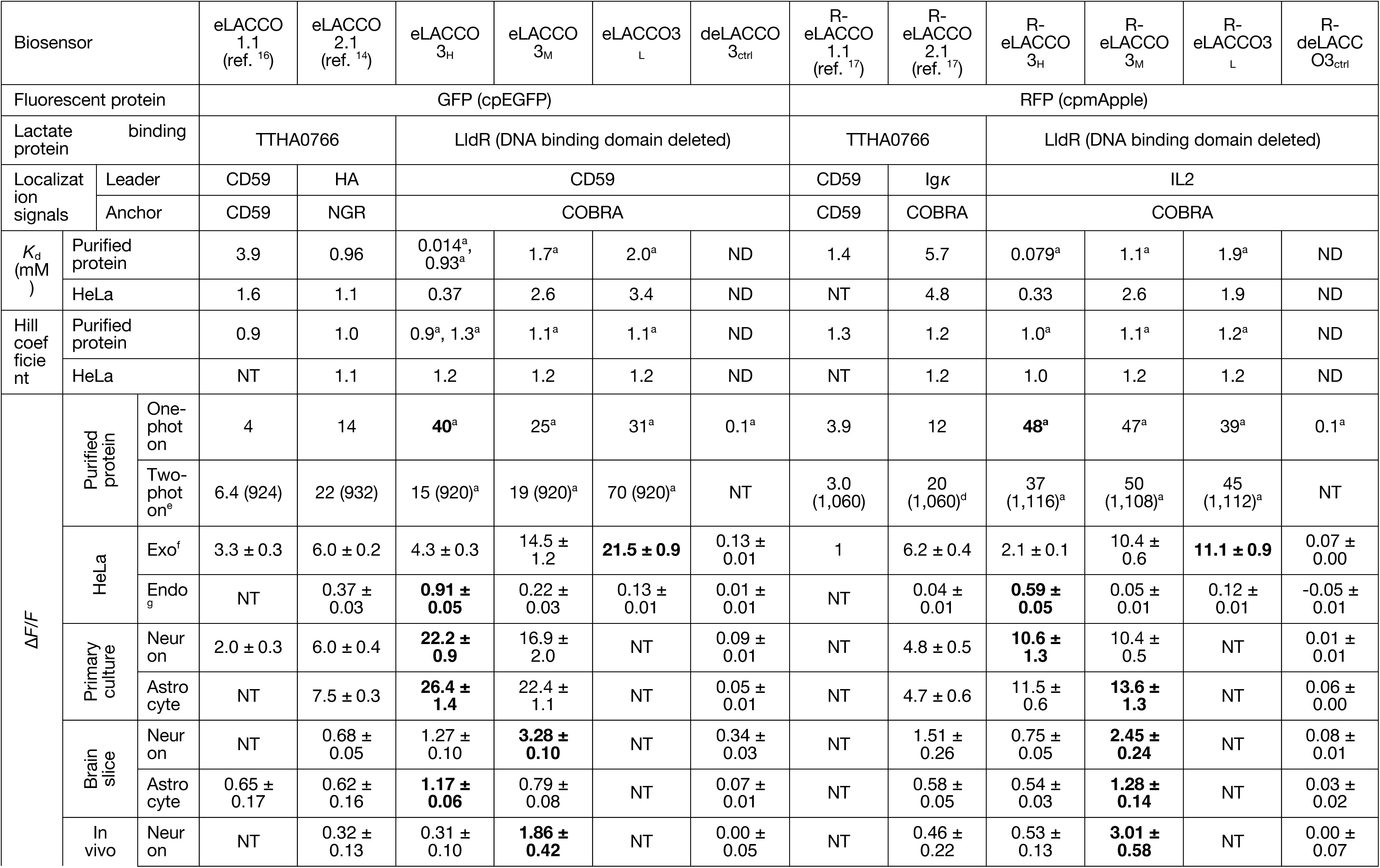
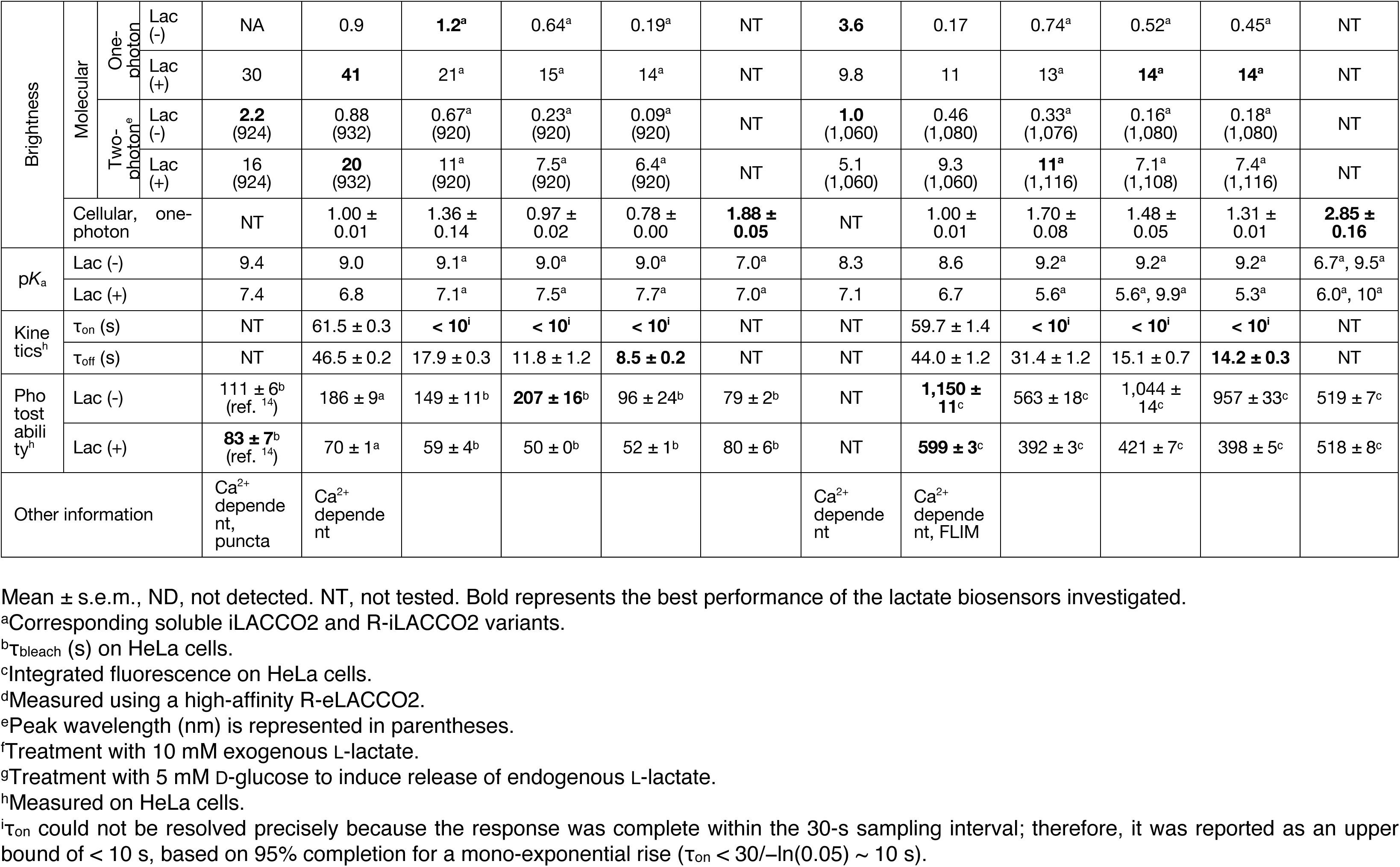
Comparison of performance for extracellular L-lactate biosensors.

### Multiplexed in vivo imaging of extracellular lactate and neuronal Ca^2+^ dynamics

In acute brain slices, electrical stimulation evoked R-eLACCO3_M_ fluorescence increases that were abolished by tetrodotoxin (TTX; **Fig. 4a–c**). The lactate-insensitive control R-deLACCO3_ctrl_ showed no detectable stimulus-evoked response (**Supplementary Fig. 8a**), supporting the detection of action potential-dependent extracellular lactate transients. To evaluate compatibility with neuronal Ca^2+^ imaging in vivo, we paired R-eLACCO3_M_ with GCaMP6f. In the olfactory bulb of anesthetized mice, R-eLACCO3_M_ was expressed on astrocyte surfaces, whereas GCaMP6f reported neuronal Ca^2+^ signals in the glomerular layer (**Fig. 4d**). A 1-s odor stimulus elicited rapid neuronal Ca^2+^ responses followed by delayed, reversible increases in R-eLACCO3_M_ fluorescence (**Fig. 4e,f**).

**Fig. 4.**
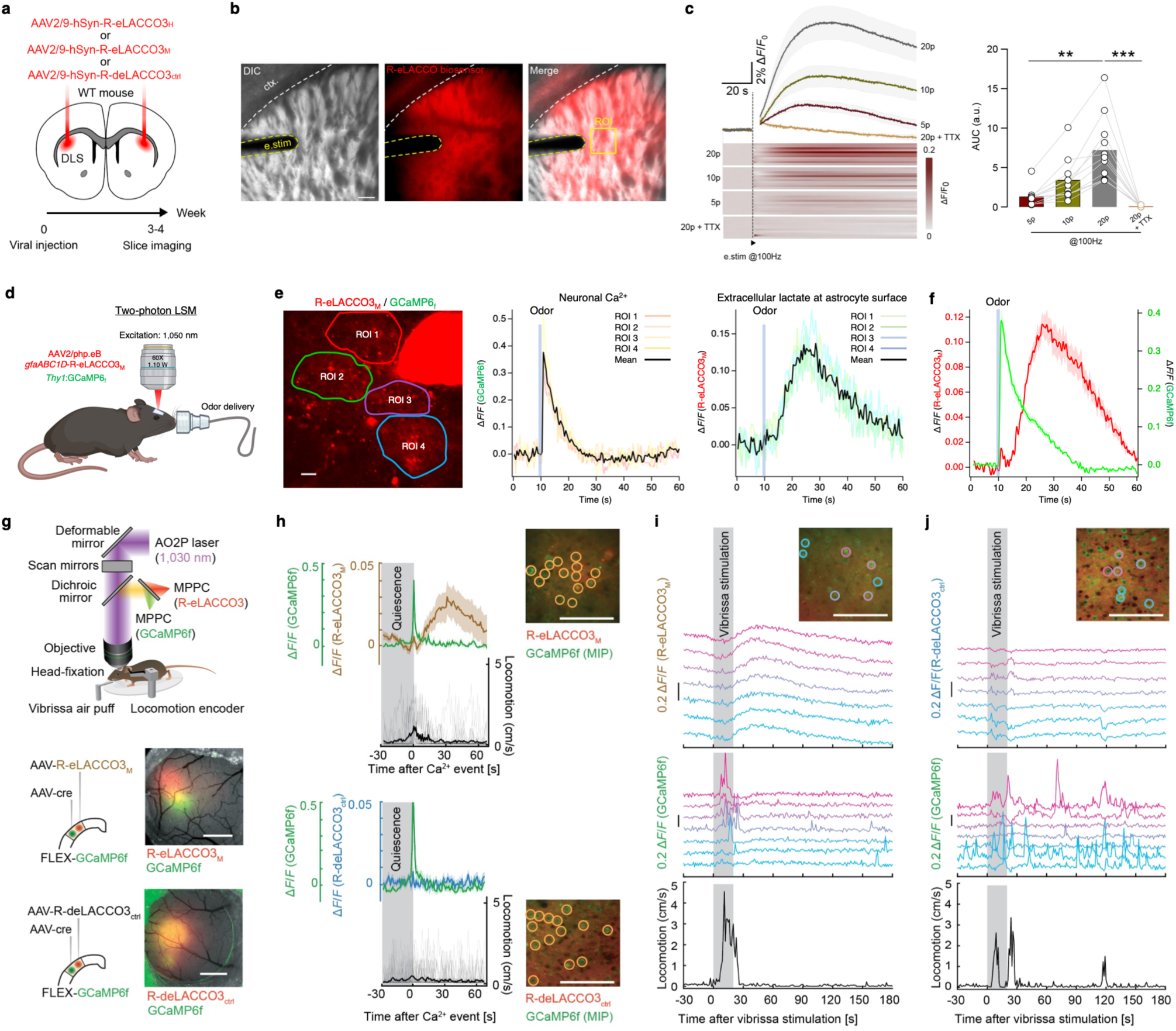
Multiplexed in vivo imaging of extracellular lactate and neuronal Ca^2+^ dynamics. (**a**) Viral expression strategy to express R-eLACCO3 variants in the dorsolateral striatum. (**b**) R-eLACCO3_M_ sensor expression from acute slice preparation. ctx, cortex; e.stim, electrical stimulation. Scale bar, 150 μm. (**c**) Fluorescence traces (left) and area under the curve (AUC) quantification (right) of R-eLACCO3_M_ following electrical stimulation applied at 100 Hz for 5,10, and 20 pulses applied at 20 s in the dorsolateral striatum. Frames containing stimulation artifact were removed. *n* = 11 slices from 4 mice. Kruskal–Wallis test followed by Friedman test. Data are presented as mean ± s.e.m. \*\**p* < 0.01, \*\*\**p* < 0.001. TTX, tetrodotoxin. (**d**) Neuronal Ca^2+^ and extracellular lactate responses were monitored in the OB glomerular layer of mice expressing YFP and GCaMP6f under the control of the M72 and Thy1 promoters, and transduced with AAV2/php.eB-gfaABC1D-R-eLACCO3_M_. All fluorophores were excited at 1,050 nm. Odors were delivered by an olfactometer. (**e**) Left: 4 astrocytes expressing R-eLACCO3_M_ and delineated by regions of interest (ROIs). The M72 glomerulus, in which olfactory sensory neuron terminals express YFP serves as a regional landmark for glomeruli sensitive to the odor ethyl tiglate. Middle: neuronal Ca^2+^ responses to odor from the 4 ROIs. Right: extracellular lactate responses measured at astrocyte surfaces from the same ROIs. Each trace represents the average of 4 stimulations. (**f**) Mean neuronal Ca^2+^ and extracellular lactate responses to odor (*n* = 27 astrocytes in 3 mice). Mean ± s.e.m. (**g**) Schematic illustration of simultaneous AO2P imaging of L-lactate and Ca^2+^ in awake mice co-expressing R-eLACCO3_M_ or R-deLACCO3_ctrl_ (red, excited at 1,030 nm) and GCaMP6f (green, excited at 1,030 nm) in vS1 neurons. Scale bars, 1 mm. (**h**) Δ*F*/*F* traces of neurons co-expressing GCaMP6f and R-eLACCO3_M_ (top, *n* = 14 neurons and 36 Ca^2+^ events after 30 seconds of quiescence) or R-deLACCO3_ctrl_ (bottom, *n* = 14 neurons and 36 Ca^2+^ events after 30 seconds of quiescence) in an awake mouse. Mean ± s.e.m. Scale bars, 100 µm. (**i**) Fluorescence response traces of R-eLACCO3_M_ and GCaMP6f in vS1 neurons of an awake mouse upon vibrissa air puff stimulation. Scale bar, 100 µm. (**j**) Fluorescence response traces of R-deLACCO3_ctrl_ and GCaMP6f in vS1 neurons of an awake mouse upon vibrissa air puff stimulation. Scale bar, 100 µm.

We next used adaptive optics two-photon (AO2P) microscopy for dual-color imaging in awake cortex. Neuronal co-expression of R-eLACCO3_M_ and GCaMP6f was achieved using hSyn-driven AAVs encoding R-eLACCO3_M_ and Cre in combination with a Cre-dependent GCaMP6f reporter (**Fig. 4g**)^21^. R-eLACCO3_M_ functionality as an extracellular L-lactate biosensor was independently validated in vivo (**Supplementary Fig. 8b–d**). In an awake mouse, recordings aligned to spontaneous neuronal Ca^2+^ events following a quiescent period showed delayed R-eLACCO3_M_ fluorescence increases; separate recordings with R-deLACCO3_ctrl_ showed little change (**Fig. 4h**). Vibrissa stimulation accompanied by locomotion was associated with R-eLACCO3_M_ fluorescence increases across multiple regions of interest (**Fig. 4i**), whereas R-deLACCO3_ctrl_ showed no comparable increases (**Fig. 4j**). Similar observations were made during spontaneous locomotion (**Supplementary Fig. 8e,f**).

Collectively, these experiments demonstrate multiplexed in vivo imaging of extracellular lactate and neuronal Ca^2+^ dynamics, with extracellular lactate signals accompanying odor-evoked neuronal activity and, in cortical recordings, spontaneous neuronal Ca^2+^ events and locomotion.

## Discussion

Here, we developed eLACCO3 and R-eLACCO3 for Ca^2+^-independent extracellular lactate imaging in complementary fluorescence colors. Their advances extend beyond larger responses to exogenous lactate: direct comparisons in cultured cells demonstrated improved detection of endogenous extracellular lactate oscillations, and applications in the mouse brain demonstrated compatibility with simultaneous measurements of neuronal Ca^2+^ signals. These capabilities support measurements of extracellular lactate in relation to brain state.

A central design choice was to replace the Ca^2+^-dependent TTHA0766 sensing scaffold used in earlier extracellular sensors^14,16,17^ with LldR-based sensor cores. Their Ca^2+^ dependence can confound lactate measurements during activity-associated extracellular Ca^2+^ fluctuations^18,22^. Our development workflow combined cysteine substitutions, linker engineering, and directed evolution of the sensor cores with screening of cell-surface targeting sequences. This approach emphasizes the importance of evaluating sensor performance in the intended cellular environment rather than assuming that properties measured for soluble proteins will transfer unchanged to the cell surface.

eLACCO3 and R-eLACCO3 outperform the previously reported extracellular L-lactate biosensors in Ca^2+^ independency, kinetics, and sensitivity, extending capabilities beyond larger responses to exogenous lactate. The high-affinity variants resolved glucose-evoked extracellular lactate oscillations around cultured HeLa cells that were less frequently detected by eLACCO2.1 or undetected by R-eLACCO2.1. Faster recovery may help distinguish successive transients that slower sensors would blur. Simultaneous intracellular imaging linked extracellular signals to oscillations within individual cells, suggesting that surface-targeted sensors report local pericellular rather than bulk-medium dynamics. R-eLACCO3_M_ also supports two-photon imaging alongside GCaMP6f, enabling comparisons between extracellular lactate and neuronal Ca^2+^ dynamics.

Two practical considerations remain. First, blue-light photoactivation of R-eLACCO3 under some cultured-cell and in vivo conditions warrants caution with green biosensors or blue-light-activated optogenetic tools. Sequential excitation and optimized illumination intensity and timing may mitigate interference; lactate-insensitive R-deLACCO3_ctrl_ should be evaluated under matched illumination to identify lactate-independent fluorescence changes. Future engineering could explore alternative red fluorescent proteins, such as mScarlet used in the Ca^2+^ indicator PinkyCaMP^23^, to minimize photoactivation. Second, sensor selection should be guided by the preparation and expected response regime. High-affinity variants performed well in glucose-stimulated cultured cells, whereas medium-affinity variants produced larger early responses in the in vivo cortical lactate-application experiments. Brightness, affinity, and kinetics should therefore be considered together rather than selecting a variant solely from its maximal Δ*F*/*F*.

In the olfactory bulb of anesthetized mice, brief odor stimulation evoked neuronal Ca^2+^ responses followed by large, reversible extracellular lactate increases. Simultaneous neuronal Ca^2+^ imaging distinguished local activity-associated responses from larger locomotion-associated elevations. These observations highlight the value of multiplexed imaging for resolving relationships to spontaneous and sensory-evoked neuronal activity. However, concentration measurements establish neither flux direction nor cellular origin, and larger locomotion-associated signals do not alone distinguish local production from broader metabolic regulation. Perturbing lactate production and transport will be necessary to test these mechanisms.

In summary, eLACCO3 and R-eLACCO3 expand the available tools for extracellular lactate imaging by combining Ca^2+^-independent responses with complementary colors and improved detection of endogenous dynamics. Their compatibility with neuronal Ca^2+^ indicators provides a framework for investigating how extracellular lactate availability relates to physiological state.

## Methods

### General methods and materials

PCR amplification was performed using Phusion high-fidelity DNA polymerase (Thermo Fisher Scientific), unless otherwise stated. Error-prone PCR was carried out using Taq DNA polymerase (New England Biolabs). Restriction enzymes, ligation kits, and GeneJET miniprep kits were obtained from Thermo Fisher Scientific. DNA fragments generated by PCR or restriction digestion were purified by agarose gel electrophoresis followed by extraction using the GeneJET gel extraction kit (Thermo Fisher Scientific). DNA sequencing was conducted by Fasmac Co., Ltd. and the Genomic Technology Core at the Institute of Plant and Microbial Biology, Academia Sinica. Fluorescence spectra were recorded using Spark plate readers (Tecan).

### Engineering of the LACCO biosensors

The green fluorescent intracellular L-lactate biosensor iLACCO1 (ref. ^15^) was first modified by introducing Cys53Ala and Cys257Ser substitutions using Q5 high-fidelity DNA polymerase (New England Biolabs), generating iLACCO1.3. This construct was subjected to iterative rounds of library generation by whole-gene error-prone PCR and screening in *E*. *coli* DH10B (Thermo Fisher Scientific) cultured in LB medium supplemented with 100 μg mL^-1^ ampicillin and 0.02% L-arabinose. In each round, ∼200–400 fluorescent colonies were selected, cultured, and analyzed in 96-well plates using a plate reader. After six rounds of screening, iLACCO1.4 was obtained. Two amino acids (Trp110 and Ser111) were subsequently inserted between the N-terminal LldR domain and cpGFP by site-directed mutagenesis. The resulting construct was further optimized through nine additional rounds of error-prone PCR, yielding iLACCO2_H_. To tune lactate affinity, Val395Thr and Val395Ile substitutions were independently introduced into iLACCO2_H_ to generate iLACCO2_M_ and iLACCO2_L_, respectively. An Asp69Asn substitution was introduced into iLACCO2_H_ to abolish lactate binding. A final round of directed evolution was performed to improve fluorescence brightness, yielding the control variant diLACCO2ctrl.

For the red fluorescent series, R-iLACCO1 (ref. ^14^) was modified by introducing Cys53Ala and Cys180Tyr substitutions using Q5 polymerase to generate R-iLACCO1.3. This variant underwent iterative rounds of whole-gene error-prone PCR and screening in *E*. *coli* using the same workflow described above. After seven rounds, R-iLACCO1.9 was identified. Mutations identified during evolution (Ala141Val, Gln154Pro, and Arg300Ser) were subsequently combined by site-directed mutagenesis to generate R-iLACCO2_H_. To modulate lactate affinity, Val391Ile and Val391Thr substitutions were independently introduced into R-iLACCO2_H_ to generate R-iLACCO2_M_ and R-iLACCO2_L_, respectively. The Asp69Asn substitution was introduced to abolish ligand binding. One additional round of directed evolution was performed to enhance brightness, yielding the control variant R-diLACCO2_ctrl_.

### Protein purification and in vitro characterization

The gene encoding iLACCO2 and R-iLACCO2 variants, with a poly-histidine tag on the N-terminus, was expressed from the pBAD vector. Bacteria were lysed with a cell disruptor (Branson) and then centrifuged at 15,000*g* for 20 min, and proteins were purified by Ni-NTA affinity chromatography (Agarose Bead Technologies). To perform pH titrations, protein solutions were diluted into buffers (pH from 4 to 11) containing 30 mM trisodium citrate, 30 mM sodium borate, 30 mM MOPS, 100 mM KCl, and either no L-lactate or 100 mM L-lactate. Fluorescence intensities as a function of pH were then fitted by a sigmoidal binding function to determine the p*K*_a_. For lactate titration, buffers were prepared by mixing a L-lactate (-) buffer (30 mM MOPS, 100 mM KCl, pH 7.2) and a L-lactate (+) buffer (30 mM MOPS, 100 mM KCl, 100 mM L-lactate, pH 7.2) to provide L-lactate concentrations ranging from 0 mM to 100 mM at 25°C. Fluorescence intensities were plotted against L-lactate concentrations and fitted by a sigmoidal binding function to determine the Hill coefficient and apparent *K*_d_.

Two-photon excitation spectra of iLACCO2 and R-iLACCO2 variants were measured according to the general methods and protocols described in refs ^24,25^. Briefly, a tunable femtosecond laser InSight DeepSee (Spectra-Physics, MKS) was used to excite fluorescence of the sample contained within a PC1 spectrofluorometer (ISS, Champaign, IL). To measure the two-photon excitation spectral shapes, we used a combination of 633SP + 680 SP + 770 SP filters for iLACCO2 variants and a combination of 633SP + 770SP filters for R-iLACCO2 variants. LDS 798 in 1:2 CHCl_3_:CDCl_3_ and Coumarin 540A in DMSO were used as spectral standards. Quadratic power dependence of fluorescence intensity in the proteins and standards was checked at several wavelengths across the spectrum. The two-photon cross section (σ_2_) of the anionic form of the chromophore in free and saturated states of both sensors was measured^25^. In the case of R-iLACCO2 variants, we used 1,060-nm for the two-photon excitation and 532 nm for the one-photon excitation. Both one-photon- and two-photon-excited fluorescence signals were measured through a combination of filters 770SP + 561 LP. In the case of iLACCO2 variants, we used 940 nm for the two-photon excitation and 488 nm for the one-photon excitation. A combination of 770SP and 520LP filters was used to collect fluorescence. Rhodamine 6G in methanol was used as a reference standard for both proteins^24^. The two-photon absorption spectra were normalized to the measured σ_2_ values. The two-photon brightness (*F*_2_) near the anionic peak was calculated by multiplying the two-photon cross section measured as described above by the quantum yield of anionic form and by the relative fraction of the anionic form of the chromophore. The whole two-photon excitation spectrum was then scaled to this *F*_2_ value to obtain the entire *F*_2_ spectrum (corresponding to the linear combination of the brightness values of the neutral and anionic forms at each excitation wavelength). The method is described in further details in refs. ^26,27^.

One-photon absorption spectra were measured using a Lambda 950 spectrophotometer (PerkinElmer). Extinction coefficients of the neutral and anionic forms and their relative fractions at neutral pH were determined by gradual alkaline denaturation of the samples containing either free or bound states^27^. Fluorescence emission and excitation spectra were measured with an LSS 55 spectrofluorimeter (PerkinElmer). Fluorescence quantum yield of anionic form was measured at several excitation wavelengths using an integration sphere instrument Quantaurus-QY (Hamamatsu). The plateau values, where the quantum yield did not depend on excitation wavelength, corresponding to excitation of only the anionic form in the red part of absorption spectrum were averaged and taken as a final value.

### Construction of mammalian expression vectors

For cytosolic expression, biosensor coding sequences were excised from pBAD vectors using XhoI and HindIII and ligated into pcDNA3.1 (Thermo Fisher Scientific). For cell surface targeting, biosensor genes were amplified by PCR, digested with BglII and EcoRI, and inserted into a modified pcDNA3.1 vector containing an N-terminal leader sequence and a C-terminal membrane anchor. Both targeting elements were derived from ref. ^17^. For AAV-based expression, biosensor coding sequences were subcloned into pAAV vectors under the control of either the hSyn or gfaABC1D promoter using standard restriction enzyme-based cloning.

### Imaging in HeLa and HEK293T cell lines

HeLa and HEK293T cells were maintained in Dulbecco’s modified Eagle medium (DMEM; Nakalai Tesque and Thermo Fisher Scientific) supplemented with 10% fetal bovine serum (FBS; Sigma-Aldrich) and 1% penicillin-streptomycin (Nakalai Tesque and Thermo Fisher Scientific) at 37 °C in a humidified incubator with 5% CO_2_. Cells were seeded onto 35-mm glass-bottom dishes (Iwaki) and transiently transfected with plasmid DNA using polyethyleneimine (Polysciences).

Live-cell imaging was performed using either an IX83 wide-field fluorescence microscope (Olympus) equipped with a pE-300 LED light source (CoolLED), a 40× oil-immersion objective (Olympus, NA 1.3), an ImagEM X2 EM-CCD camera (Hamamatsu), CellSens software (Olympus), and a stage-top incubator (Tokai Hit), or an Axio Observer 7 wide-field fluorescence microscope (Zeiss) equipped with a Viluma 7 LED light source (Zeiss), a 40× oil-immersion objective (Zeiss, NA 1.4), an ORCA-Fusion BT CMOS camera (Hamamatsu), ZEN software (Zeiss), and a stage-top incubator (Tokai Hit). For green fluorescent biosensors (iLACCOs, eLACCOs, EGFP), filter sets consisted of excitation 470/20 nm, a 490-nm dichroic mirror, and emission 518/45 nm. For red fluorescent biosensors (R-iLACCOs, R-eLACCOs, mApple, mCherry), excitation 545/20 nm, a 565-nm dichroic mirror, and emission 598/55 nm were used. Image analysis was performed using ImageJ (National Institutes of Health).

For membrane localization analysis, HEK293T cells were co-transfected with GFP-based eLACCO biosensor constructs together with pDisplay-mCherry (Ig*κ*–mCherry–PDGFR) or with RFP-based R-eLACCO biosensors together with pAMEXT-EGFP (CD59–EGFP–CD59), which served as plasma membrane markers. Fluorescence profiles from the biosensor and membrane marker channels were combined, and Pearson correlation coefficients were calculated using the *corr* function in the Pandas package (v1.5.3). For quantification of basal brightness and Δ*F*/*F*, HeLa cells were transfected with the indicated LACCO biosensor constructs. Basal fluorescence intensity was measured within a cellular region of interest prior to L-lactate addition and defined as brightness. Δ*F*/*F* was calculated by comparing fluorescence intensities before and after application of 10 mM L-lactate.

For photostability measurements, transfected HeLa cells were continuously illuminated at full LED intensity (∼10 mW cm^-2^ for iLACCO/eLACCO variants and ∼4 mW cm^-2^ for R-iLACCO/R-eLACCO variants, measured at the objective). Time-lapse images were acquired for 4 min with an exposure time of 50 ms and no interval between frames. For L-lactate bath application to measure in situ *K*_d_ and response kinetics, HeLa cells grown on coverslips were transfected with plasmids encoding eLACCO/R-eLACCO variants. At 48 h post-transfection, coverslips were mounted in Attofluor^TM^ cell chambers (Thermo Fisher Scientific) containing Hank’s balanced salt solution (HBSS(+); Nakalai Tesque and Thermo Fisher Scientific) supplemented with 10 mM HEPES and 1 μM AR-C155858 (Tocris). Bath solution exchange during imaging was performed in a remove-and-add manner using a homemade solution remover^28^. For Ca^2+^-dependent fluorescence imaging, transfected cells were transferred to Attofluor^TM^ chambers containing HBSS(−) supplemented with 10 mM HEPES and 10 mM L-lactate 48 h after transfection. Solutions supplemented with Ca^2+^ of 100 nM, 1 μM, 10 μM, 100 μM, 300 μM, 1 mM, 3 mM, and 10 mM were applied. Buffer exchange was performed as described above.

For basal fluorescence intensity measurements, HeLa cells expressing the indicated lactate biosensors under unstimulated conditions were detached and analyzed using a BD FACSymphony A1. Basal fluorescence intensity was measured for each sample. Cells expressing iLACCO or eLACCO variants were excited with a 488-nm laser, and emission was collected using a 530/30 nm bandpass filter. Cells expressing R-iLACCO or R-eLACCO variants were excited with a 561-nm laser, and fluorescence was detected using a 610/20 nm bandpass filter. The geometric mean fluorescence intensity was calculated for each samples. Flow cytometry data were analyzed using FlowJo software version 10.10.0.

### Fluorescence imaging of primary neurons and astrocytes

Primary cortical and hippocampal cultures were prepared from embryonic day 21 (E21) Sprague Dawley rat pups (Charles River Laboratories; purchased from Japan SLC, Inc.) obtained from a single timed-pregnant dam. Embryos were collected by Caesarean section, and dissociated cells were plated onto glass-bottom 24-well plates (Cellvis) at a density of 0.5 × 10^6^ cells per three wells. Cells were nucleofected at the time of plating using the Nucleofector 4D system (Lonza), and imaging was performed 14 days post-plating. For each construct, three wells were prepared and analyzed. Neurons were maintained in NbActive4 medium (BrainBits) supplemented with 1% penicillin-streptomycin at 37 °C in a humidified incubator with 5% CO_2_. Astrocytes were cultured in DMEM supplemented with 10% FBS and 1% penicillin-streptomycin under the same conditions. For imaging, culture medium was replaced with 1 mL of imaging buffer containing 145 mM NaCl, 2.5 mM KCl, 10 mM D-glucose, 10 mM HEPES, 2 mM CaCl_2_, and 1 mM MgCl_2_ (pH 7.4).

### Packaging and purification of adeno-associated viruses (AAVs)

AAVs were produced in HEK293T17 cells by triple transfection with the helper plasmid pXX680, a plasmid encoding AAV2/5 or AAV2/9 rep/cap, and a pAAV vector carrying the LACCO biosensor gene. Cells were harvested 48 h post-transfection by gentle scraping and pelleted by centrifugation. Viral particles were released by four freeze–thaw cycles using a dry ice–ethanol bath, followed by treatment with benzonase to degrade residual nucleic acids. AAVs were purified by iodixanol step-gradient ultracentrifugation (15%, 25%, 40%, and 60%) at 67,000 rpm for 70 min at 16 °C. The virus-containing fraction was collected, washed, and concentrated using a 100 kDa molecular weight cutoff Amicon filter. During the final concentration step, the buffer was exchanged to 5% D-sorbitol and 0.001% pluronic acid for storage and subsequent use. Viral genome titers were determined by TaqMan droplet digital PCR (ddPCR) using primers targeting the AAV2 inverted terminal repeat (ITR).

### Viral injections and acute slice preparation for sensor validation

All surgical procedures were conducted under general anesthesia using continuous isoflurane (induction at 5%, maintenance at 1.5–2% vol/vol, Fujifilm, 099-06571). Following induction of anesthesia, mice were fitted into a stereotaxic frame (David Kopf Instruments) with their heads secured by ear bars and their noses placed into a veterinary-grade anesthesia and ventilation system (VetEquip; 911103). The surgical incision site was then cleaned with 70% ethanol (vol/vol). Craniotomies (1– 2 mm in diameter) were made powered by a high-speed drill at the specific coordinates based on the position of the bregma. Viral injections were performed using beveled glass pipettes (World Precision Instruments, 1B100-4) lowered into the orbitofrontal cortex (OFC; AP: +2.6 mm, ML: 1.3 mm, DV: 2.2 mm), and 0.8 μL of AAV was delivered at 0.1 μL min^-1^ using a syringe pump (World Precision Instruments, SmartTouch Pump). The glass pipette was withdrawn 5 min after the infusion to prevent the backflow. Mice were then sutured, returned to their home cage, and allowed to rest for three weeks before acute slice imaging was conducted. For simultaneous imaging of eLACCO and R-eLACCO signals ex vivo, the AAV combinations listed in the table below were co-injected.

| eLACCO | R-eLACCO |
| --- | --- |
| AAV2/5-gfaABC1D-eLACCO2.1 | AAV2/9-hSyn-R-eLACCO2.1 |
| AAV2/5-gfaABC1D-eLACCO3<sub>H</sub> | AAV2/9-hSyn-R-eLACCO3<sub>H</sub> |
| AAV2/5-gfaABC1D-eLACCO3<sub>M</sub> | AAV2/9-hSyn-R-eLACCO3<sub>M</sub> |
| AAV2/5-gfaABC1D-deLACCO3<sub>ctrl</sub> | AAV2/9-hSyn-R-deLACCO3<sub>ctrl</sub> |
| AAV2/9-hSyn-eLACCO2.1 | AAV2/5-gfaABC1D-R-eLACCO2.1 |
| AAV2/9-hSyn-eLACCO3<sub>H</sub> | AAV2/5-gfaABC1D-R-eLACCO3<sub>H</sub> |
| AAV2/9-hSyn-eLACCO3<sub>M</sub> | AAV2/5-gfaABC1D-R-eLACCO3<sub>M</sub> |
| AAV2/9-hSyn-deLACCO3<sub>ctrl</sub> | AAV2/5-gfaABC1D-R-deLACCO3<sub>ctrl</sub> |

Mice were deeply anesthetized with isoflurane and decapitated with sharp shears. The brains were placed and coronally sliced in ice-cold modified artificial cerebrospinal fluid (aCSF) containing the following (in mM): 194 sucrose, 30 NaCl, 4.5 KCl, 1 MgCl_2_, 26 NaHCO_3_, 1.2 NaH_2_PO_4_, and 10 D-glucose, saturated with 95% O_2_ and 5% CO_2_. A vibratome (NeoLinear Slicer MT, Dosaka EM, Japan) was used to cut 300 µm brain sections. The slices were allowed to equilibrate for 30 min at 32– 34 °C in normal aCSF containing (in mM); 124 NaCl, 4.5 KCl, 2 CaCl_2_, 1 MgCl_2_, 26 NaHCO_3_, 1.2 NaH_2_PO_4_, and 10 D-glucose continuously bubbled with 95% O_2_ and 5% CO_2_. Slices were then stored at 21–23 °C in the same buffer until use.

### Ex vivo lactate imaging in acute slices for sensor validation

Individual slices were transferred to a submerged recording chamber (room temperature). All brain slices were continuously perfused at room temperature using a peristaltic pump (MP-2, Gilson Inc.). Initially, slices were perfused with recording buffer (RB) for 5 minutes, followed by a 20-minute perfusion with RB containing 1 µM AR-C155858 (HY-13248, MedChemExpress) and 1 µM VB124 (S9929, Selleck Chemicals) to inhibit monocarboxylate transporters (referred to as MCTi-RB). Subsequently, the perfusion solution was sequentially switched to MCTi-RB supplemented with sodium L-lactate (L7022-10G, Sigma-Aldrich) at concentrations of 1 and 10 mM, with each condition maintained for 10 minutes. Ex vivo imaging was conducted using a two-photon microscope (A1R MP, NIKON) equipped with a femtosecond laser system (Alcor Dual, Spark Lasers) and a water-immersion objective lens (16×, NA 0.80, NIKON). The excitation laser wavelengths for eLACCO and R-eLACCO sensors were 920 nm or 1,064 nm, respectively. Images (512 × 512 pixels, 0.78 µm/pixel) were acquired at a frame rate of ∼0.2 Hz by sequentially applying 920 nm and 1,064 nm lasers for 1 second each with 1 second intervals. Image acquisition was performed using NIS-Elements AR (version 5.30.05, NIKON). The mean fluorescence intensity was calculated over the entire field of view (FOV), and eLACCO and R-eLACCO responses were quantified as Δ*F*/*F*_0_, where *F*_0_ was defined as the mean intensity during the 100 seconds prior to lactate application.

### In utero electroporation for in vivo imaging of mouse pups

In utero electroporation was performed as described^20^ with minor modifications. Pregnant mice were anesthetized at embryonic day 15 (E15) with a combination of sodium pentobarbital (50 mg kg^-1^ body weight) and isoflurane (1.0–1.5% in air). A midline laparotomy was conducted to expose the uterus, and the DNA solution containing pK031 TRE-Cre (5 ng μL^-1^, Addgene 69136) and CAG-loxP-STOP-loxP-eLACCO-IRES-tTA-WPRE or CAG-loxP-STOP-loxP-R-eLACCO-IRES-tTA-WPRE (1 μg μL^-1^) was injected into the lateral ventricle of each embryo using a pulled glass capillary. Square electric pulses (1-3 pulses, 40 V, 50 ms, 1 Hz) were delivered using an electroporator and forceps-type electrodes (BEX, Japan). After electroporation, the uterus was repositioned, and the abdominal wall and skin were sutured. Mice were allowed to recover on a heating pad maintained at 37 °C.

### In vivo lactate imaging in mouse pups

Craniotomies for intravital imaging were performed as previously described^29^ with some modifications. Pups were anesthetized with isoflurane (0.7–1.5%), and the scalp was removed using surgical scissors. The skull overlying the cerebral cortex was then carefully removed using a 27-gauge needle while leaving the dura intact. A custom-made titanium bar (approximately 30 mg) was attached adjacent to the cranial window using dental cement. Throughout the surgical procedure, the exposed dura was continuously kept moist with cortex buffer (125 mM NaCl, 5 mM KCl, 10 mM D-glucose, 10 mM HEPES, 2 mM CaCl_2_, 2 mM MgSO_4_, pH 7.4)^30^.

In vivo imaging was performed using an SP8MP multiphoton microscope (Leica) equipped with an Alcor 920 nm-2 W laser (Spark Lasers) for eLACCO imaging, a Fidelity-2 1,070 nm laser (Coherent) for R-eLACCO imaging, and a 25× objective lens (0.95 NA). Fluorescence emission was collected through a 500–550 nm and 575–620 nm bandpass filters for eLACCO and R-eLACCO, respectively. Imaging was performed under isoflurane anesthesia (0.7–1.5%). Time-lapse images of layer 2/3 neurons were acquired at 2–4 Hz with a resolution of 512 × 512 pixels. After baseline imaging, lactate was added directly to the immersion cortex buffer to achieve a final concentration of 10 mM, and imaging was continued.

Time-lapse images were analyzed using FIJI (ImageJ, NIH). Regions of interest (ROIs) were drawn around the soma of individual neurons, and the mean fluorescence intensity of each ROI was measured over time. Baseline fluorescence (*F*_0_) was defined as the mean fluorescence intensity during the 1-min baseline period. Changes in fluorescence were calculated as Δ*F*/*F*_0_ = (*F* − *F*0)/*F*_0_, where *F* is the fluorescence intensity at each time point. For quantitative analysis, the mean Δ*F*/*F*_0_ during the 30–60 s period following lactate application was calculated for each neuron.

### Viral injections and acute slice preparation for electrical stimulation

C57BL/6J wildtype mice from both sex were used at 8 to 16 weeks of age. Mice were group-housed and maintained on a 12 h light/dark cycle (lights on 07:00–19:00) with ad libitum access to food and water. All experiments were performed during the light phase of the cycle. Acute brain slices for imaging were obtained from 8–16-week-old mice. Mice were culled by cervical dislocation within ∼2 hrs after start of light ON period of light cycle and brains were dissected out and submerged in ice-cold cutting solution containing (in mM): 194 sucrose, 30 NaCl, 4.5 KCl, 1 MgCl_2_, 26 NaHCO_3_, 1.2 NaH_2_PO_4_, and 10 D-glucose. Coronal slices 300 μm-thick containing striatum were prepared from dissected brain tissue using a vibratome (VT1200S, Leica Microsystems) and transferred to a holding chamber containing aCSF containing (in mM): 130 NaCl, 2.5 KCl, 26 NaHCO_3_, 2.5 CaCl_2_, 2 MgCl_2_, 1.25 NaH_2_PO_4_, and 10 D-glucose. Sections were incubated at 34 °C for 15 min before they were stored at room temperature (20–22 °C) in a physiological glucose aCSF at 2.5 mM until recordings were performed. All recordings were obtained within ∼8 h of slicing. All solutions were saturated with 95% O_2_ and 5% CO_2_. Before recording, individual slices were hemisected and transferred to a recording chamber and superfused at ∼2.5–3.0 mL min^-1^ with aCSF at ∼32–33 °C.

Mice were anesthetized with 4% isoflurane (O_2_ flow of ∼1 L min^-1^) and subsequently maintained with ∼1.5–2% isoflurane during surgery. Mice were placed in a small animal stereotaxic frame (David Kopf 124 Instruments), and body temperature was maintained at ∼37 °C. After exposing the skull under aseptic techniques, a small burr hole was drilled and AAV solutions were injected at an infusion rate of 200 nL min^-1^ with a 32-gauge Hamilton syringe (Hamilton Company) using a microsyringe pump (World Precision Instruments) and withdrawn 5 min after the end of injection. To virally express the sensor, the viral construct was injected bilaterally into the dorsal striatum (AP + 0.8 mm, ML ± 1.8 mm from bregma, DV − 2.40 mm from exposed dura mater) of 8–12-week-old C57BL/6J wildtype mice. Mice were used for experiments 3–4 weeks post-intracranial injection.

### Ex vivo lactate imaging in acute slices by electrical stimulation

An Olympus BX51WI microscope equipped with a 575 nm CoolLED light pE-300ultra, Iris 9 Scientific CMOS camera (Teledyne Photometrics), 629 ± 35.5 nm emission filter (Thorlabs), and 10×/0.3NA water-immersion objective (Olympus) was used for wide-field fluorescence imaging of R-eLACCO3 variants in striatal slices. Image acquisition was controlled using Micro-Manager 1.4. Electrical stimulation, LED light, and image acquisition were synchronized using TTL-driven stimuli via Multi Channel Stimulus II (Multi Channel Systems). Image files were analyzed with MATLAB R2025a and Fiji.

For experiments imaging extracellular lactate levels in response to trains of electrical stimulation, images were acquired at 10 Hz (exposure duration, 100 ms) during continuous green light (575 nm; 1 mW mm^-2^; CoolLED light pE-300ultra) for a 2 min recording window. For evoked lactate release, electrical stimulus pulses (5, 10, and 20 pulses at 100 Hz; pulse duration, 200 μs; 0.6 mA) were given by a local bipolar concentric Pt/Ir electrode (inner diameter, 25 μm; outer diameter, 125 μm; FHC) at the 20 s time point.

For analysis, we extracted fluorescence intensity from the ROI (200 × 200 μm) ∼100 μm from the stimulating electrode and derived a background-subtracted fluorescence intensity (*F*_t_) by subtracting background fluorescence intensity from an equal-sized ROI where there was no R-eLACCO3 expression (i.e., cortex). Data are expressed as a change in fluorescence (Δ*F*/*F*_0_) and were derived by calculating [(*F*_t_ – *F*_0_)/*F*_0_], where *F*_0_ is the average Ft of the first initial 20 s baseline preceding electrical stimulation.

### Surgery and AAV injection for olfactory bulb (OB) imaging

Mice were fed ad libitum and housed in a 12-hour light-dark cycle at 22 °C and 50% humidity. A total of three Thy1::GCaMP6f+/-; M72-IRES-ChR2-YFP+/+ double transgenic mice were used in this experiment. Both males and females, aged 5 to 7 months, were included. Thy1::GCaMP6f (GP5.11)^31^ mice were obtained from Jackson Laboratory and crossed with M72-IRES-ChR2-YFP^32^ kindly provided by Thomas Bozza (Department of Neurobiology, Northwestern University, USA). Permanent cranial windows were implanted over the dorsal OB as previously described^33^. Briefly, mice were anaesthetised with an intraperitoneal (IP) bolus injection of a mixture of ketamine and medetomidine (100 mg kg^-1^ and 0.5 mg kg^-1^ body weight, respectively). If necessary, an additional 10–20% of the same mixture was administered to maintain the depth of anaesthesia. Hypoxaemia was prevented by oxygen supplementation. Body temperature was monitored using a rectal probe and maintained at 36.5 °C with a feedback-controlled heating pad. A craniotomy was carefully performed using a dental drill while cold HEPES-buffered artificial cerebrospinal fluid (125 mM NaCl, 5 mM KCl, 2 mM CaCl_2_, 2 mM MgSO_4_, 10 mM D-glucose, and 10 mM HEPES, pH 7.4) was continuously applied to prevent overheating and tissue damage during the procedure. A manually trimmed coverslip (100 µm thick) was used as a cranial window and sealed with photopolymerizable dental cement. Finally, a titanium head bar was secured to the skull with dental cement. Mice were allowed to recover for at least one week before injection of the AAV vector encoding the biosensor.

Mice implanted with a cranial window were briefly anaesthetised with isoflurane (4% induction, 2.5% maintenance) and retro-orbitally injected with 3 × 10^11^ genome copies (GCs) of AAV2/PHP.eB-gfaABC1D-R-eLACCO3_M_. This vector encodes R-eLACCO3_M_ under the control of the short astrocytic promoter gfaABC1D and is packaged in an AAV2/PHP.eB capsid, which enables efficient transduction across the blood–brain barrier following systemic delivery and exhibits broad central nervous system (CNS) tropism, particularly towards neurons and astrocytes in mice^34^. The vector was diluted in PBS containing 0.001% Pluronic F-68.

### Two-photon laser scanning microscopy (TPLSM) for OB imaging

In vivo two-photon imaging of glomerular astrocytes was carried out in mice anaesthetised with a mixture of ketamine and medetomidine (100 mg kg^-1^ and 0.5 mg kg^-1^, respectively). Experiments were performed within 20–120 minutes following the injection of anaesthetic. Animal breathed regularly at 2–4 Hz and their body temperature was maintained at approximately 36.5 °C using a heating pad.

A home-built olfactometer based on the design of the Rinberg lab (Janelia olfactometer design) and controlled by custom LabVIEW software was used to deliver the odor (ethyl tiglate). Air was continuously delivered to the mouse nose at 500 mL min^-1^ and, upon stimulation, a valve switched the flow from air to the odour-air mixture for the duration of the stimulation (1 s). The flows of the air and odor lines were balanced such that no pressure changes occurred during the switch. An additional oxygen flow of 50 mL min^-1^ was continuously delivered through the nose cone to increase the oxygen concentration of the air or odor–air mixture reaching the nose of the animal to 29%. Ethyl tiglate was delivered at 22.5%.

TPLSM imaging was performed using a custom-built microscope^35^, and data were acquired using customised LabVIEW software (National Instruments). Two-photon excitation was achieved using a femtosecond laser from Spectra Physics (Insight, 70 fs pulses, 80 MHz). Laser power was modulated with an acousto-optic modulator (AA Optoelectronic, MT110B50-A1.5-IR-Hk). Galvanometric mirrors (Cambridge Technology) were used to direct the beam to the desired locations within the sample. The excitation beam was focused onto the sample using a LUMFLN60XW water-immersion objective (1.10 NA, Olympus). R-eLACCO3M and GCaMP6f were excited at 1,050 nm, and the emitted fluorescence was separated from the excitation light with a dichroic mirror (DXCR 875, Chroma Technology Corp.). Emitted fluorescence was then separated into green and red channels using a dichroic mirror (cut-off wavelength: 560 nm). The green (GCaMP6f) channel was filtered using low-pass and band-pass filters (E800 and HQ 525/50 nm, Chroma Technology Corp.), and fluorescence was collected with a GaAsP photomultiplier tube (Hamamatsu). The red (R-eLACCO3_M_) channel was filtered using a low-pass filter (E800, Chroma) and a band-pass filter (FF01-650/150, Semrock), and fluorescence was collected with a red-sensitive photomultiplier tube (R6357, Hamamatsu). Time-series images were acquired at 2-2.5 Hz.

GCaMP6f and R-eLACCO3_M_ fluorescence signals were extracted from selected ROIs using customised LabVIEW software (National Instruments) and were computed as Δ*F*/*F* = (*F* – *F*_0_)/*F*_0_, where *F*0 represents the baseline fluorescence over the first 10 s before stimulation, and *F* represents the fluorescence at time *t*. Fluorescence signals from each astrocyte-associated ROI were first averaged per mouse, and the plotted data represent the mean across mice ± standard deviation. Individual ROI responses (R-eLACCO3_M_ and GCaMP6f) are average of several stimulations. To enable temporal alignment for averaging recordings with slightly different sampling rates, Δ*F*/*F* traces were first interpolated to a uniform temporal grid (0.1 s resolution) in MATLAB (using the “pchip” function, R2022b, MathWorks).

### Surgery and AAV injection for adaptive optics two-photon (AO2P) microscopy

Awake imaging was performed in male and female mice aged 4–9 months. Mice were bred from C57BL/6J animals (Jackson Laboratory) and maintained at 20–26 °C and 30–70% humidity under a 12 h light–dark cycle with food and water available ad libitum. Surgical procedures were performed under isoflurane anesthesia (4% in oxygen for induction and 1–2% for maintenance). Buprenorphine was administered subcutaneously for analgesia (0.1 μg g^-1^ body weight), and body temperature was maintained at 36–37 °C using a heating pad. A cranial window and viral injections were prepared over the right vibrissal primary somatosensory cortex (vS1; center, 1.5 mm posterior to bregma and 3.4 mm lateral to the midline), following a previously described AO2P preparation with modifications^17^. A 3.5-mm craniotomy was made while preserving the dura. Glass injection pipettes (Drummond, QF100-60-10) were pulled using a P-2000 puller (Sutter Instruments, P-2000) and trimmed to a 10–20 μm outer diameter. AAVs were delivered at multiple sites in vS1, 400 μm below the dura, using a motorized micromanipulator (Sutter Instruments, MPC-200, MP285/M-21826, and ROE-200) and syringe pump (KD Scientific). For R-eLACCO3_M_ imaging, 50–150 nL AAV2/9-hSyn-R-eLACCO3_M_ (1.1 × 10^12^ GCs mL^-1^) was injected. For simultaneous lactate and Ca^2+^ imaging, AAV2/9-hSyn-R-eLACCO3M was mixed with AAV8-hSyn-Cre-WPRE-hGH (Addgene, 105553-AAV8) and injected into heterozygous GCaMP6f reporter mice (Jackson Laboratory, 028865). The cranial window consisted of a 3.5-mm round coverslip (Warner Instruments, no. 1 CS-3.5R, 64-0739) attached to a 5-mm round coverslip (Warner Instruments, no. 1 CS-5R, 64-0700) using Norland Optical Adhesive 61. The window was placed into the craniotomy, sealed with cyanoacrylate glue (Loctite, 401), and reinforced with Metabond (Parkell L-Powder and Quick Base). A custom titanium head bar was fixed to the skull with Metabond, and exposed bone was covered with dental acrylic (Lang Ortho-Jet). Imaging was performed 1–6 months after virus injection following at least three 1-h habituation sessions for head fixation.

### AO2P microscopy

AO2P imaging was conducted in awake head-fixed mice using adaptive correction with a deformable mirror to compensate microscope-induced aberrations^17^. Excitation was provided by a Chameleon Discovery NX femtosecond pulsed laser (Coherent) tuned to 1,030 nm for simultaneous imaging of R-eLACCO3 and GCaMP6f. Images were acquired at 1 Hz using a resonant–galvo scanner with a 25× objective lens (Olympus, XLPlanN), dichroic mirrors (FF775-Di01 and FF662-Di01), emission filters (FF01-593/46 and FF01-240-sp), and MPPC detectors (Hamamatsu, 20A-002). Post-objective laser power was kept below 40 mW for all imaging. The dura was aligned to the imaging plane using two goniometers (GNL10/M) under epifluorescence guidance (Zeiss Stereo, X-Cite 120Q). For intravenous infusion experiments, tail-vein catheters (custom-made 30G needle connected to PE10 tubing) were placed under 1.5% isoflurane after warming the tail at 37 °C for 5 min. Catheter placement was verified by injection of 50 μL saline. After recovery from anesthesia, saline and L-lactate (Sigma, L7022) were infused through PE10 tubing connected to calibrated syringe pumps via Y-connectors (Instechlab, SCY-25). Image acquisition, locomotion, and vibrissa stimulation were tracked using LabChart (ADInstruments) and analyzed using Matlab (MathWorks). Translational transformation was used to register each frame to a smoothed, averaged frame at the start of each trial for motion correction, and frames with excess translation vector (>1 s.d.) were excluded and interpolated. Ca^2+^ events larger than 4 s.d. were considered significant.

### Data representation and statistical analysis

Data are presented as mean ± s.d. or mean ± s.e.m., as indicated in the figure legends. Box plots show the median (center line), 25th and 75th percentiles (box limits), and whiskers extending to ±1 s.d. from the mean (black or white filled circle). Sample sizes (*n*) are indicated in the corresponding figures or legends. Statistical analyses were performed, as indicated in the figure legends, using GraphPad Prism 9. Data were plotted using Microsoft Excel and Igor Pro 8. Differences were considered significant at \**p* < 0.05, \*\**p* < 0.01, \*\*\**p* < 0.001, or \*\*\*\**p* < 0.0001 and not significant at *p* > 0.05.

## Ethical statement

All experiments performed at University of California San Diego have been approved by the Institutional Animal Care and Use Committee at University of California, San Diego. All procedures involving animals used for acute slice experiments were approved by the RIKEN Animal Care and Use Committee. All animal care and experimentations at Institut de la Vision were performed in accordance with the INSERM Animal Care and Use Committee guidelines and approved by the ethical committee (protocol number 27135 2020091012114621). For experiments performed at The University of Tokyo, all methods for animal care and use were approved by the institutional review committees of the School of Science, The University of Tokyo. All animal experiments performed at Kumamoto University have been approved by the Institutional Animal Care and Use Committee at Kumamoto University. All procedures at University of Oxford were performed in accordance with the Animals in Scientific Procedures Act 1986 (UK) (Amended 2012) with ethical approval from an Animal Welfare and Ethical Review Body at the University of Oxford, under the authority of Licenses PP8860348 from the UK Home Office.

## Data availability

The plasmids generated in this study have been deposited in Addgene.

## Acknowledgments

The authors thank Academia Sinica IBC Biophysics Core Facility and IPMB Genomic Technology Core for technical support. Work at Academia Sinica (AS) was supported by AS (AS-CDA-115-L01 and AS-GCP-115-L03), National Science and Technology Council (NSTC, 115-2113-M-001-009-MY3), and the Japan Science and Technology Agency (JST, JPMJPR22E9 and JPMJCR24T6). Work at Montana State University was supported by the NIH/NINDS BRAIN grant U24 NS109107. Work at RIKEN was supported by Japan Society for the Promotion of Science (JSPS, JP24K02128, JP24H02180, and JP23H04179), JST (JPMJFR2249), and Japan Agency for Medical Research and Development (AMED, JP22dk0207063 and JP23jf0126004). Work at the University of Oxford was supported by the Medical Research Council (UKRI533) to S.J.C, Aligning Science Across Parkinson’s (ASAP-025192) to S.J.C and by a postdoctoral EMBO fellowship (ALTF 1572-2025) to T.B.

## Author contributions

Y.N., J.-J.L., and Y.K. developed the LACCO biosensors and performed in vitro characterization. M.D. measured absorption and two-photon excitation spectra. Y.N. and Y.K. carried out screening of leader and anchor sequences. T.N. performed flow cytometry analysis. Q.-L.T. and M.B. prepared plasmids for bioresource distribution. M.B. and M.-E.P. produced AAVs. H.K. and J.N. performed ex vivo characterization. C.-Y.W. and H.M. carried out in vivo validation. T.B. and S.J.C. performed electrostimulation experiments. E.T.-T and S.C. performed OB imaging. P.M. and D.K. conducted AO2P imaging. Y.N. supervised the study and wrote the manuscript.

---

**Supplementary Fig. 1.**
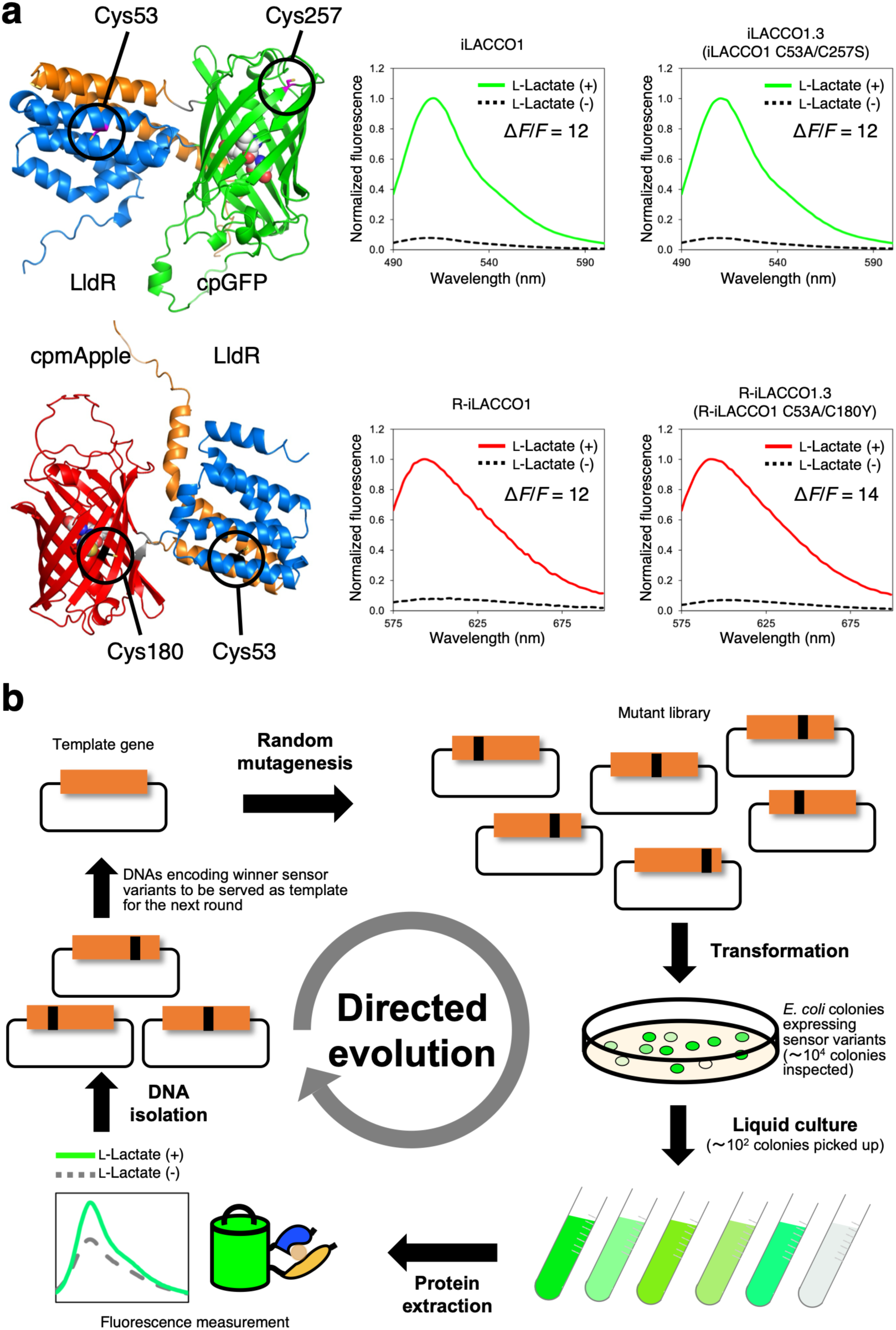
Engineering of iLACCO and R-iLACCO variants. (**a**) Cysteine substitutions introduced into iLACCO1 (top) and R-iLACCO1 (bottom). AlphaFold structural models (left) indicate the positions of cysteine residues. Fluorescence emission spectra of crude protein extracts (right) were measured in the presence (10 mM) or absence of L-lactate. (**b**) Schematic of the directed evolution workflow for iLACCO variants. Either selected regions (for example, linker sequences) or the full-length gene of the template L-lactate biosensor were subjected to random mutagenesis, and the resulting libraries were expressed in *E*. *coli*. Fluorescent colonies were selected, cultured, and lysed for screening based on Δ*F*/*F* in response to 10 mM L-lactate. Variants exhibiting the largest responses were pooled and used as templates for subsequent rounds. The same workflow was applied to R-iLACCO variants.

**Supplementary Fig. 2.**
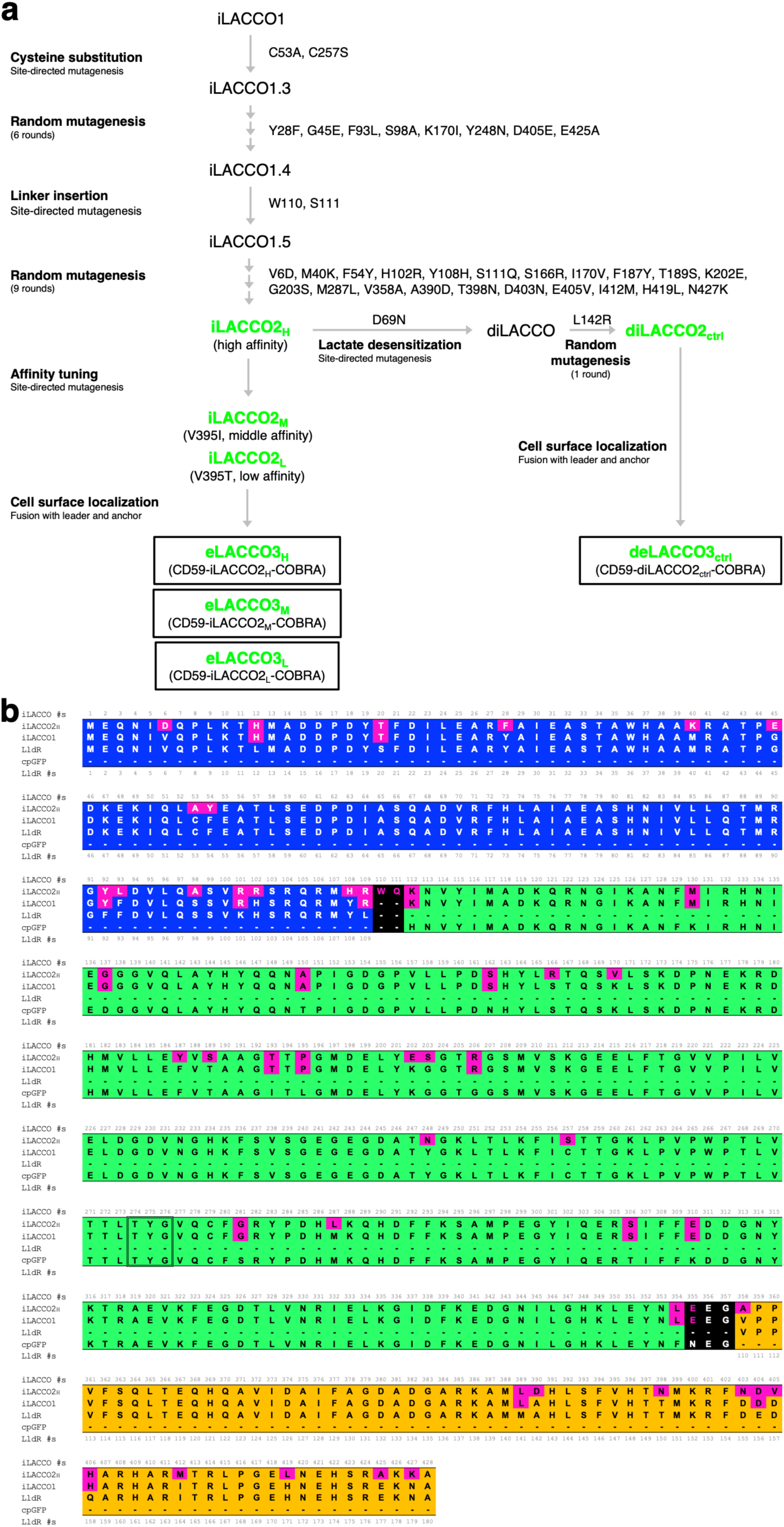
Lineage and sequence of iLACCO2 variants. (**a**) Lineage of LACCO variants. (**b**) Sequence alignment of iLACCO2H, iLACCO1, LldR, and cpGFP. Mutations in iLACCO variants, relative to LldR and cpGFP, are highlighted in magenta. The chromophore-forming residues are surrounded by double line.

**Supplementary Fig. 3.**
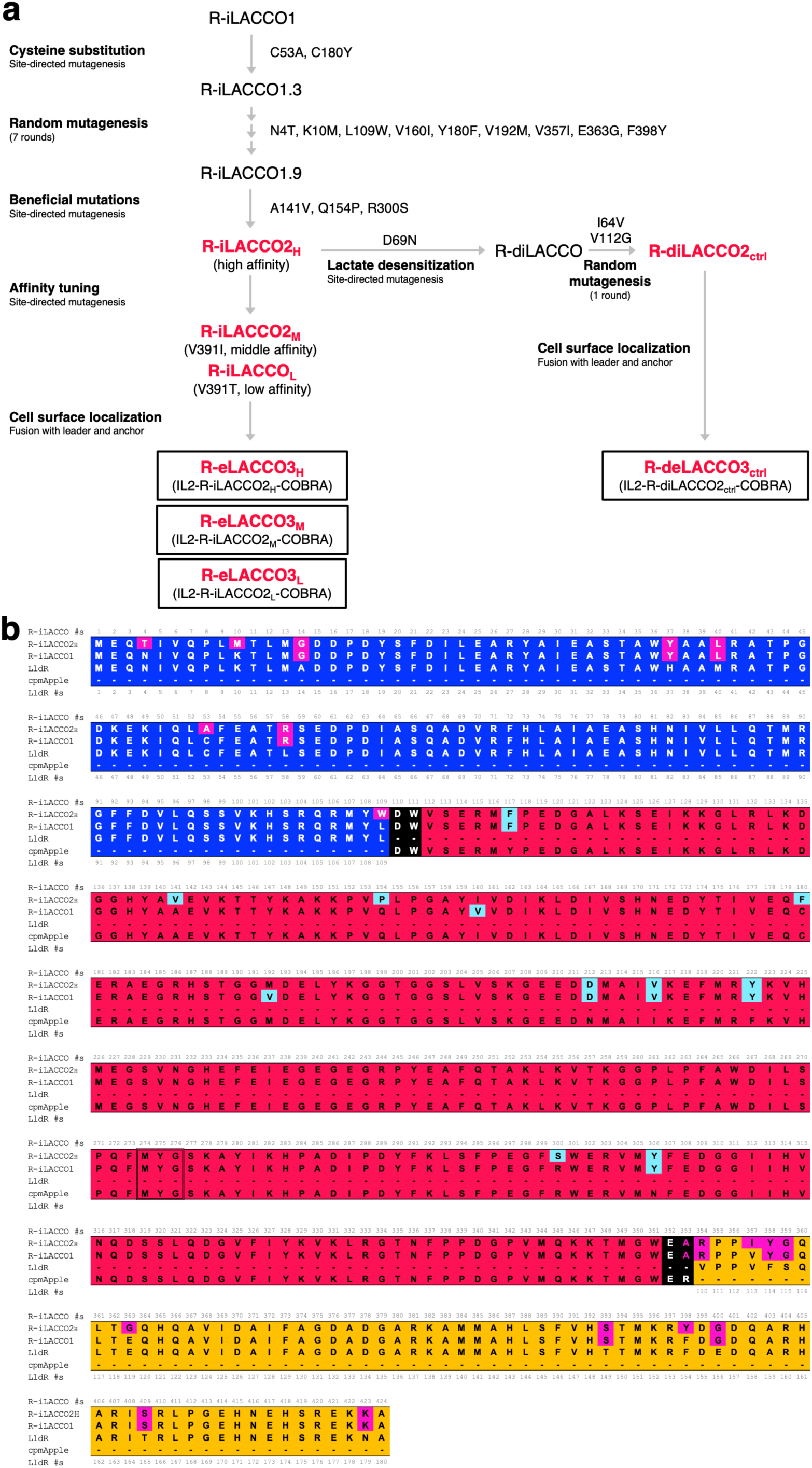
Lineage and sequence of R-iLACCO2 variants. (**a**) Lineage of R-LACCO variants. (**b**) Sequence alignment of R-iLACCO2_H_, R-iLACCO1, LldR, and cpmApple. Mutations in R-iLACCO variants, relative to LldR and cpmApple, are highlighted in magenta and light blue, respectively. The chromophore-forming residues are surrounded by double line.

**Supplementary Fig. 4.**
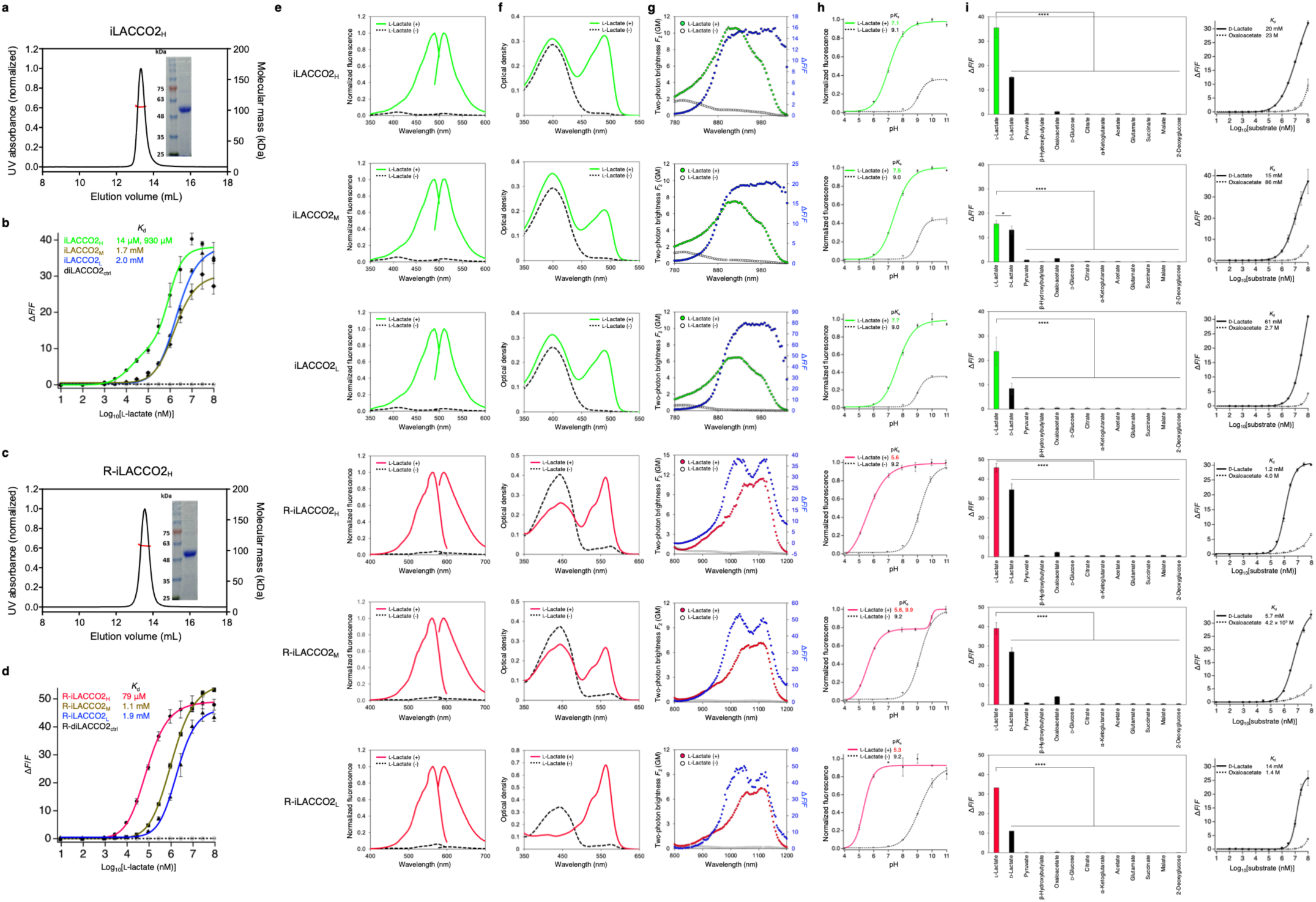
In vitro characterization of iLACCO2 and R-iLACCO2 variants. (**a**) Size-exclusion chromatography and SDS-PAGE profile of iLACCO2_H_ protein. (**b**) Dose-response curves of iLACCO variants for L-lactate. *n* = 3 experimental triplicates (mean ± s.d.). (**c**) Size-exclusion chromatography and SDS-PAGE profile of R-iLACCO2_H_ protein. (**d**) Dose-response curves of R-iLACCO variants for L-lactate. *n* = 3 experimental triplicates (mean ± s.d.). (**e**) Excitation and emission spectra in the presence (10 mM) and absence of L-lactate. (**f**) Absorbance spectra in the presence (10 mM) and absence of L-lactate. (**g**) Two-photon excitation spectra in the presence (10 mM) and absence of L-lactate. GM, Goeppert-Mayer units. (**h**) pH titration curves in the presence (100 mM) and absence of L-lactate. *n* = 3 experimental triplicates (mean ± s.d.). (**i**) Pharmacological specificity. Concentration of each metabolite is 10 mM. *n* = 3 experimental triplicates (mean ± s.d.). Right figures show dose-response curves for D-lactate and oxaloacetate. *n* = 3 experimental triplicates (mean ± s.e.m.). Statistical analyses were performed using one-way ANOVA with the Dunnett’s post hoc tests. \**p* = 0.0458, \*\*\*\**p* < 0.0001.

**Supplementary Fig. 5.**
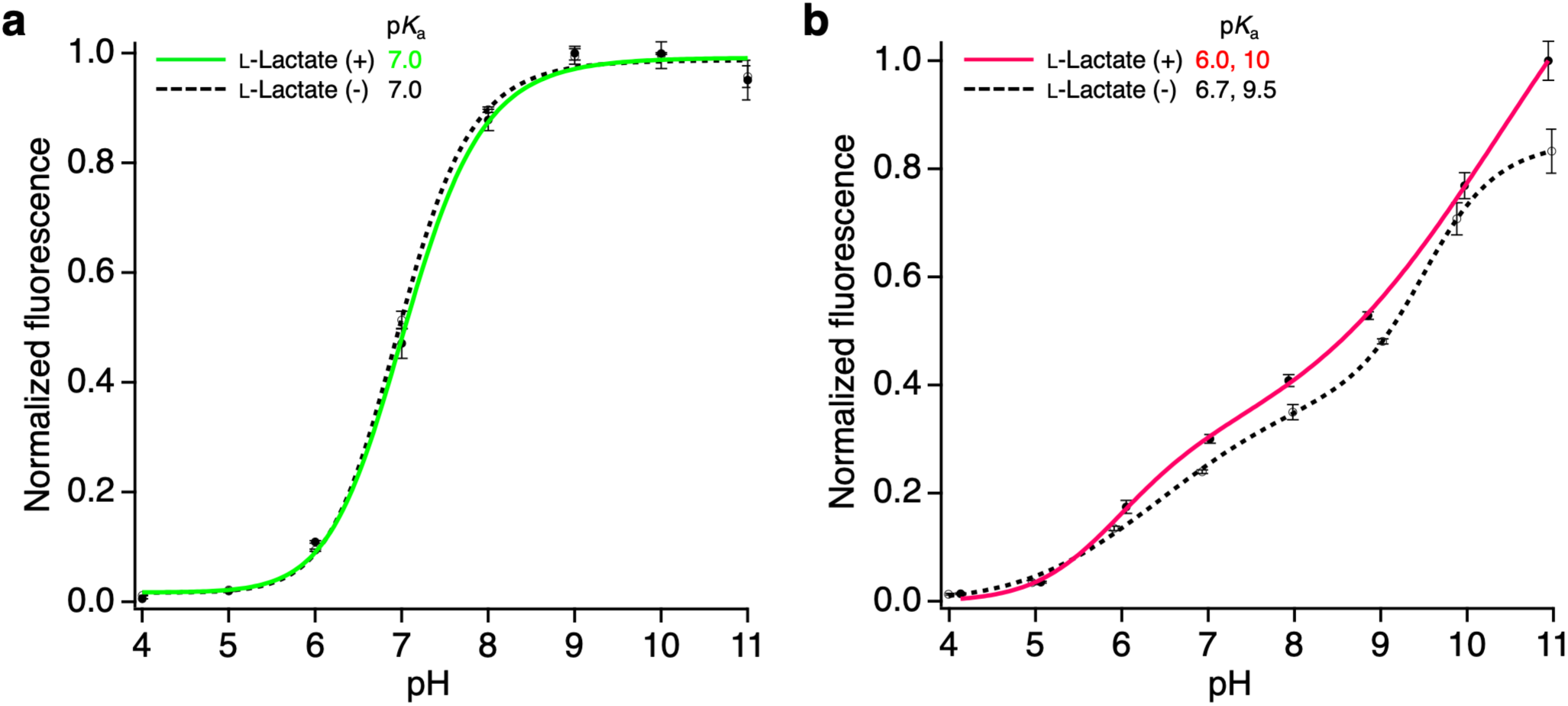
In vitro characterization of diLACCO2_ctrl_ and R-diLACCO2_ctrl_. (**a**) pH titration curves of diLACCO2ctrl in the presence (100 mM) and absence of L-lactate. *n* = 3 experimental triplicates (mean ± s.d.). (**b**) pH titration curves of R-diLACCO2ctrl in the presence (100 mM) and absence of L-lactate. *n* = 3 experimental triplicates (mean ± s.d.).

**Supplementary Fig. 6.**
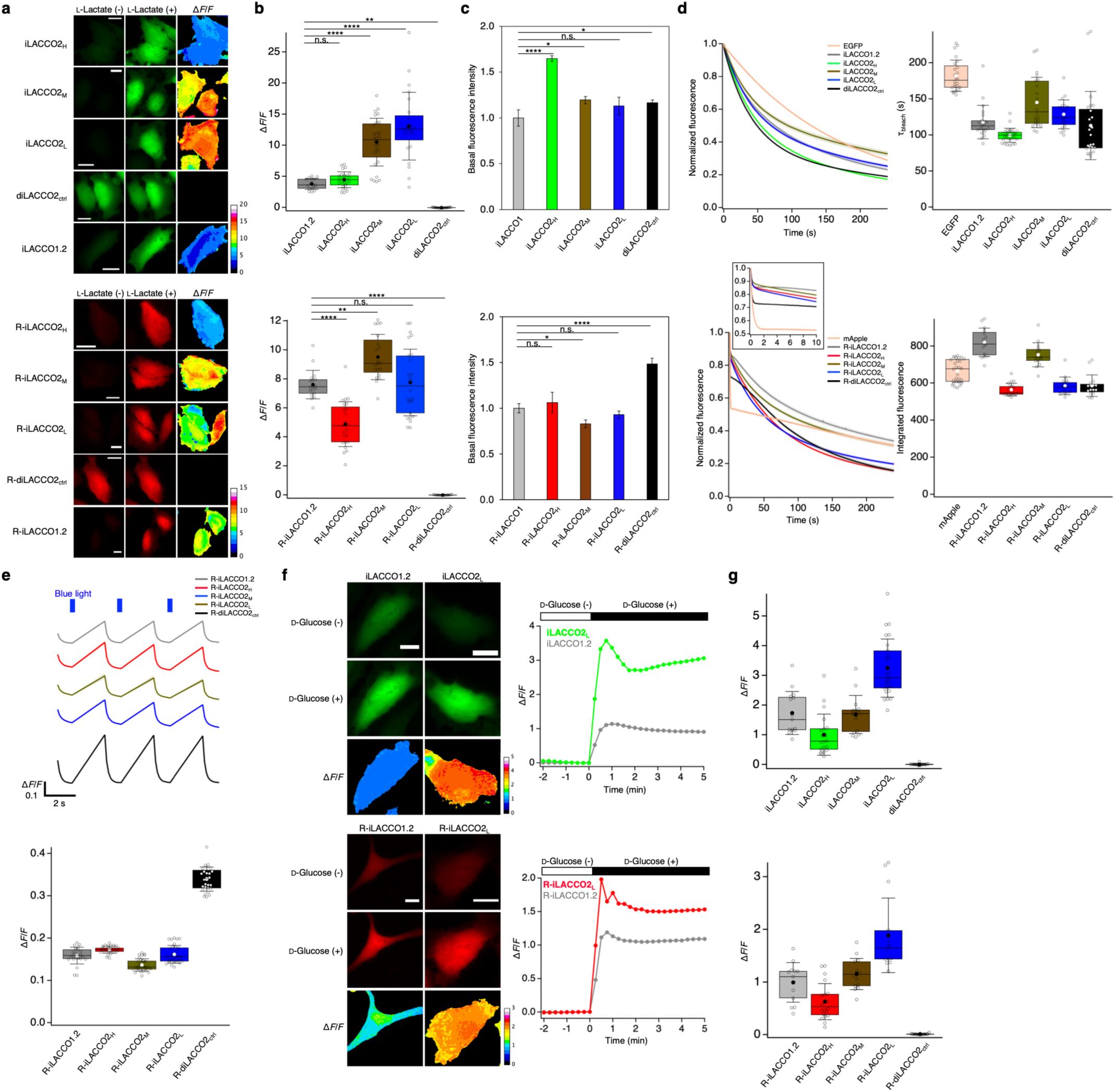
Live cell characterization of iLACCO and R-iLACCO variants. (**a**) Fluorescence images in HeLa cells expressing iLACCO (top) and R-iLACCO (bottom) biosensors. Scale bars, 10 μm. (**b**) Δ*F*/*F* in HeLa cells in (a). *n* = 17, 26, 28, 24, 22, 18, 24, 20, 26, and 17 cells for iLACCO1.2, iLACCO2_H_, iLACCO2_M_, iLACCO2_L_, diLACCO2_ctrl_, R-iLACCO1.2, R-iLACCO2_H_, R-iLACCO2_M_, R-iLACCO2_L_, and R-diLACCO2_ctrl_, respectively. One-way ANOVA with the Dunnett’s post hoc tests. *p* (iLACCO1.2 vs. iLACCO2_H_) = 0.8793, \*\**p* (iLACCO1.2 vs. diLACCO2_ctrl_) = 0.0010, \*\**p* (R-iLACCO1.2 vs. R-iLACCO2_M_) = 0.0013, *p* (R-iLACCO1.2 vs. R-iLACCO2_L_) = 0.9946, \*\*\*\**p* < 0.0001. (**c**) Basal fluorescence intensity of iLACCO (top) and R-iLACCO (bottom) variants expressed in HeLa cells. *n* = 3 experimental triplicates (mean ± s.e.m.). One-way ANOVA with the Dunnett’s post hoc tests. \*\*\*\**p* (iLACCO1 vs. iLACCO2_H_) < 0.0001, \**p* (iLACCO1 vs. iLACCO2_M_) = 0.0131, *p* (iLACCO1 vs. iLACCO2_L_) = 0.0924, \**p* (iLACCO1 vs. diLACCO2_ctrl_) = 0.0335, *p* (R-iLACCO1 vs. R-iLACCO2_H_) = 0.6325, \**p* (R-iLACCO1 vs. R-iLACCO2_M_) = 0.0324, *p* (R-iLACCO1 vs. R-iLACCO2_L_) = 0.05317, \*\*\*\**p* (R-iLACCO1 vs. R-diLACCO2_ctrl_) < 0.0001. (**d**) Photobleaching curves (left, mean ± s.e.m.) and τ_bleach_ or integrated fluorescence (right) of iLACCO (top) and R-iLACCO (bottom) variants expressed in HeLa cells using one-photon wide-field microscopy (∼10 mW cm^-2^ at 470 nm for iLACCO and ∼4 mW cm^-2^ at 560 nm for R-iLACCO). *n* = 24, 21, 29, 22, 19, 24, 28, 15, 19, 15, 17, and 16 cells for EGFP, iLACCO1.2, iLACCO2_H_, iLACCO2_M_, iLACCO2_L_, diLACCO2ctrl, mApple, R-iLACCO1.2, R-iLACCO2_H_, R-iLACCO2_M_, R-iLACCO2_L_, and R-diLACCO2_ctrl_, respectively. The inset shows the photobleaching curves of R-iLACCO variants in the first 10 seconds. (**e**) Fluorescence traces (top, mean ± s.e.m.) and Δ*F*/*F* (bottom) in response to blue light (460–480 nm) in HeLa cells expressing R-iLACCO variants. *n* = 24, 31, 32, 32, and 28 cells for R-iLACCO1.2, R-iLACCO2_H_, R-iLACCO2_M_, R-iLACCO2_L_, and R-diLACCO2_ctrl_, respectively. (**f**) Fluorescence images (left) and traces (right) in HeLa cells expressing iLACCO (top) and R-iLACCO (bottom) variants upon 5 mM D-glucose treatment after starvation. Scale bars, 10 μm. (**g**) Δ*F*/*F* in HeLa cells expressing iLACCO (top) and R-iLACCO (bottom) variants upon 5 mM D-glucose treatment after starvation. *n* = 15, 20, 15, 25, 16, 13, 20, 14, 14, and 17 cells for iLACCO1.2, iLACCO2_H_, iLACCO2_M_, iLACCO2_L_, diLACCO2_ctrl_, R-iLACCO1.2, R-iLACCO2H, R-iLACCO2_M_, R-iLACCO2L, and R-diLACCO2ctrl, respectively.

**Supplementary Fig. 7.**
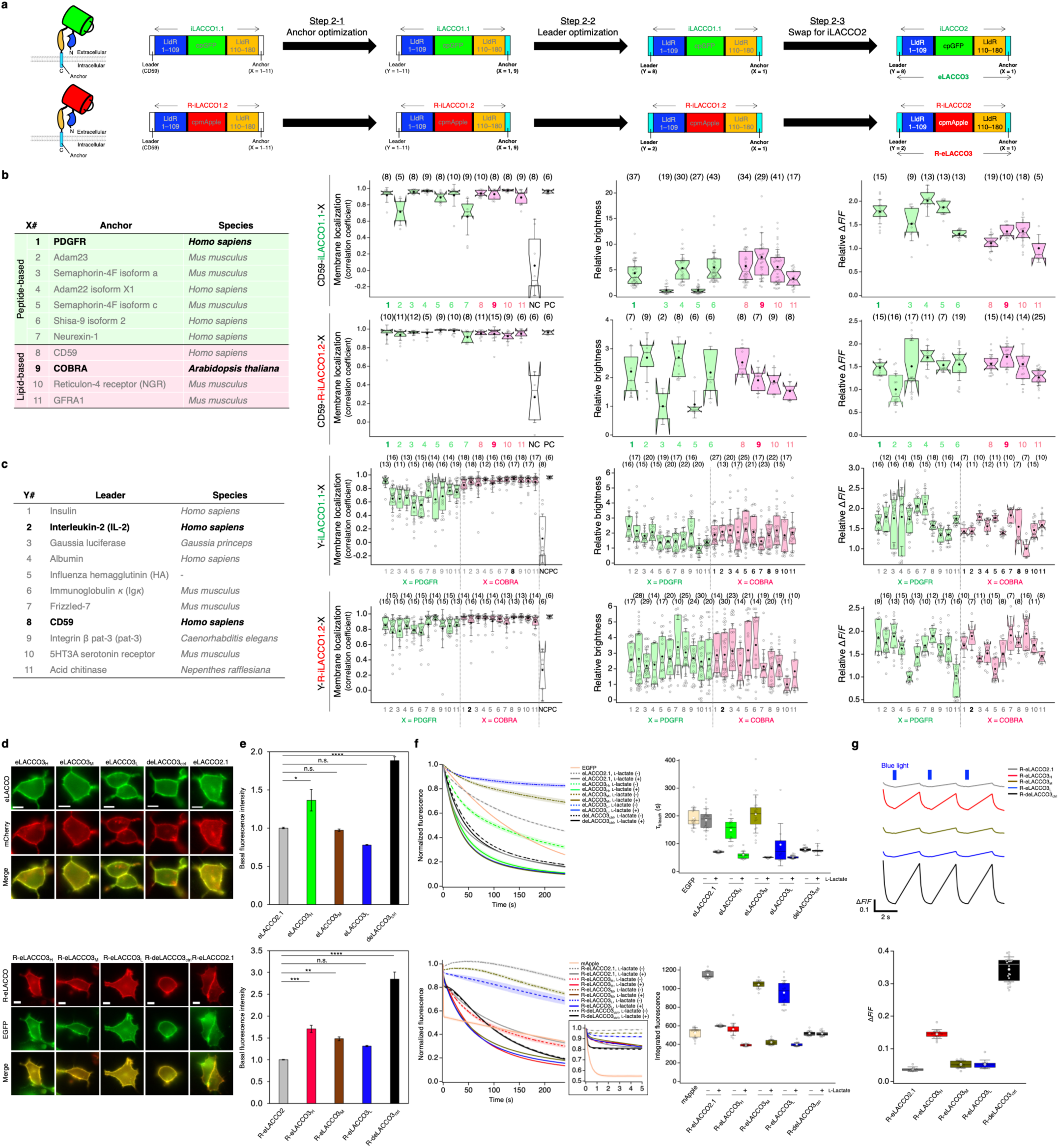
Optimization of leader/anchor combination and live cell characterization of eLACCO and R-eLACCO variants. (**a**) Overview of the leader/anchor optimization. (**b**) Membrane localization, relative brightness, and relative Δ*F*/*F* of iLACCO1.1 and R-iLACCO1.2 with CD59 leader and a range of anchors in HEK293T (membrane localization) and HeLa cells (relative brightness and Δ*F*/*F*). Parentheses represent number of cells investigated. (**c**) Membrane localization, relative brightness, and relative Δ*F*/*F* of iLACCO1.1 and R-iLACCO1.2 with various leader sequences and PDGFR or COBRA GPI anchor in HEK293T (membrane localization) and HeLa cells (relative brightness and Δ*F*/*F*). Parentheses represent number of cells investigated. NC represents negative control in which cells express cytosolic mCherry for iLACCO1.1, and cytosolic EGFP for R-iLACCO1.2. PC represents positive control in which cells express Ig*κ*-mCherry-PDGFR for iLACCO1.1, and CD59-EGFP-CD59 for R-iLACCO1.2. (**d**) Localization of eLACCO and R-eLACCO variants in HEK293T cells. mCherry with Ig*κ* leader and PDGFR anchor and EGFP with CD59 leader and CD59 anchor were used as cell surface marker. Scale bars, 10 μm. (**e**) Basal fluorescence intensity of eLACCO (top) and R-eLACCO (bottom) variants expressed on HeLa cells. *n* = 3 experimental triplicates (mean ± s.e.m.). One-way ANOVA with the Dunnett’s post hoc tests. \**p* (eLACCO2.1 vs. eLACCO3_H_) = 0.0112, *p* (eLACCO2.1 vs. eLACCO3_M_) = 0.9947, *p* (eLACCO2.1 vs. eLACCO3_L_) = 0.1294, \*\*\*\**p* (eLACCO2.1 vs. deLACCO3_ctrl_) < 0.0001, \*\*\**p* (R-eLACCO2.1 vs. R-eLACCO3_H_) = 0.0006, \*\**p* (R-eLACCO2.1 vs. R-eLACCO3_M_) = 0.0098, *p* (R-eLACCO2.1 vs. R-eLACCO3_L_) = 0.0989, \*\*\*\**p* (R-eLACCO2.1 vs. R-deLACCO3_ctrl_) < 0.0001. (**f**) Photobleaching curves (left, mean ± s.e.m.) and τ_bleach_ or integrated fluorescence (right) of eLACCO (top) and R-eLACCO (bottom) variants in the presence (10 mM) and absence of L-lactate on HeLa cells. *n* = 19, 19, 18, 15, 15, 19, 18, 10, 22, 19, 13, 16, 14, 15, 13, 15, 11, 16, 16, 17, 14, and 13 cells for EGFP, eLACCO2.1 (L-lactate -/+), eLACCO3_H_ (L-lactate -/+), eLACCO3_M_ (L-lactate -/+), eLACCO3_L_ (L-lactate -/+), deLACCO3_ctrl_ (L-lactate -/+), mApple, R-eLACCO2.1 (L-lactate -/+), R-eLACCO3H (L-lactate -/+), R-eLACCO3_M_ (L-lactate -/+), R-eLACCO3_L_ (L-lactate -/+), and R-deLACCO3_ctrl_ (L-lactate -/+), respectively. (**g**) Fluorescence traces (mean ± s.e.m.) and Δ*F*/*F* in response to blue light on HeLa cells expressing R-eLACCO variants. *n* = 8, 16, 14, 12, and 20 cells for R-eLACCO2.1, R-eLACCO3_H_, R-eLACCO3_M_, R-eLACCO3_L_, and R-deLACCO3_ctrl_, respectively.

**Supplementary Fig. 8.**
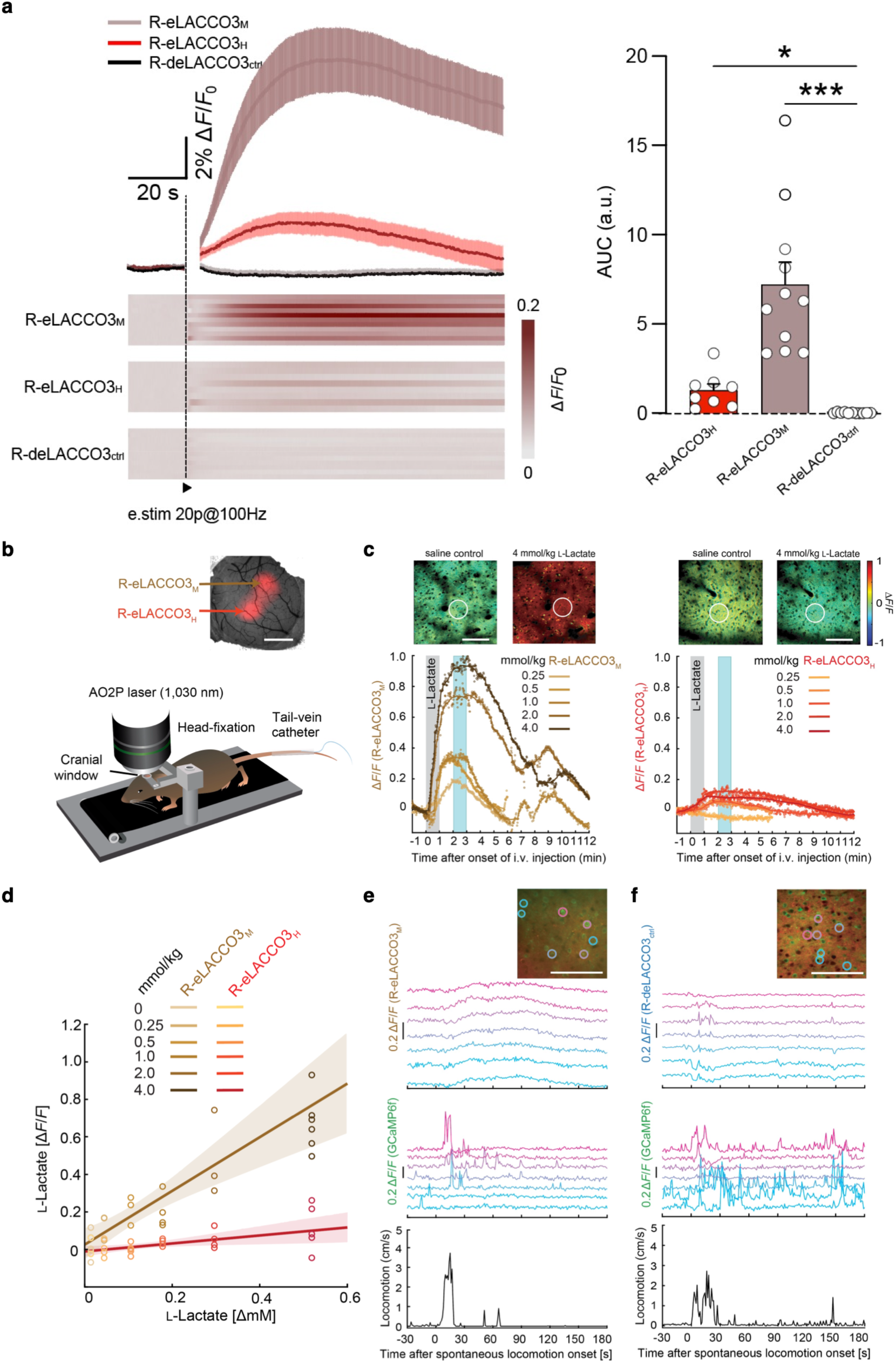
Ex vivo and in vivo L-lactate imaging using R-eLACCO3 variants. (**a**) Fluorescence traces (left) and AUC quantification (right) of R-eLACCO3 variants following electrical stimulation (20p @100 Hz) applied at 20 s in the dorsolateral striatum. Frames containing stimulation artifact were removed. *n* = 8-11 slices from 4 mice. Kruskal–Wallis test followed by Dunn’s post-hoc multiple-comparison test. Data are presented as mean ± s.e.m. \**p* < 0.05, \*\*\**p* < 0.001. (**b**) Schematic illustration of in vivo AO2P imaging for awake head-restrained mice expressing R-eLACCO3_H_ and R-eLACCO3_M_ in neurons of primary vibrissa somatosensory (vS1) cortex during tail vein injections of L-lactate. Scale bar, 1 mm. (**c**) Signal traces of R-eLACCO3_M_ (left) and R-eLACCO3_H_ (right) using AO2P with increasing dosages of L-lactate injection. The ratiometric images (top) show the ROI (white circle) used for the traces. Scale bars, 100 μm. (**d**) Correlation plot between the signals of R-eLACCO3_M_ (3 mice) or R-eLACCO3H (3 mice) and electrochemical probe (x axis) in awake mice upon i.v. injection of L-lactate. Slope is 0.2 for R-eLACCO3_H_ (*R*^2^ = 0.24) and 1.4 for R-eLACCO3_M_ (*R*^2^ = 0.62). (**e**) Fluorescence response traces of R-eLACCO3_M_ and GCaMP6f in vS1 neurons of an awake mouse in a spontaneous locomotion event (starting at time point 0) after 30 seconds of rest. Scale bar, 100 μm. (**f**) Fluorescence response traces of R-deLACCO3_ctrl_ and GCaMP6f in vS1 neurons of an awake mouse in a spontaneous locomotion event (starting at time point 0) after 30 seconds of rest. Scale bar, 100 μm.

**Supplementary Note 1 | In vitro characterization of iLACCO2 and R-iLACCO2 variants.**

Purified iLACCO2_H_ exhibited one- and two-photon Δ*F*/*F* values of 40 and 15, respectively, and biphasic binding with apparent *K*_d_ values of 14 and 930 μM. R-iLACCO2_H_ showed corresponding responses of 48 and 37 and monophasic binding (*K*_d_ = 79 μM; **Supplementary Fig. 4a–e**). Mutating Val395 in iLACCO2 or Val391 in R-iLACCO2 generated medium- and low-affinity variants with millimolar *K*_d_ values (**Supplementary Fig. 4b,d**). H, M, and L designations reflect the relative lactate affinities of the corresponding purified sensor cores; their apparent affinities can differ after cell-surface targeting (**Fig. 1d**). Asp69Asn abolished lactate binding; subsequent brightness optimization yielded diLACCO2_ctrl_ and R-diLACCO2_ctrl_ (**Supplementary Figs. 2,3**). Characterization of absorbance spectra, two-photon excitation spectra, pH titration, and molecular specificity is provided in **Supplementary Figs. 4f–l** and **5**.

In HeLa cells, lactate elicited Δ*F*/*F* values of 4.4 ± 0.3, 10.5 ± 0.7, and 13.1 ± 1.1 for iLACCO2_H_, iLACCO2_M_ and iLACCO2_L_, respectively, versus 3.8 ± 0.2 for iLACCO1.2 (mean ± s.e.m.). Corresponding R-iLACCO2_H_, R-iLACCO2_M_ and R-iLACCO2_L_ responses were 4.9 ± 0.3, 9.5 ± 0.3, and 7.7 ± 0.5, versus 7.6 ± 0.2 for R-iLACCO1.2; controls were unresponsive (**Supplementary Fig. 6a,b**). Flow cytometry showed generally greater brightness for iLACCO2 than iLACCO1; R-iLACCO2_H_ exceeded R-iLACCO1, whereas R-iLACCO2_M_ and R-iLACCO2_L_ showed lower fluorescence (**Supplementary Fig. 6c**). iLACCO2_M_ and iLACCO2_L_ were more photostable than iLACCO1.2, whereas iLACCO2_H_ was less stable. Integrated fluorescence, used because photobleaching of mApple-based sensors was non-exponential^37^, identified R-iLACCO2_M_ as the most photostable engineered variant, although R-iLACCO1.2 remained superior (**Supplementary Fig. 6d**). Red variants showed modest blue-light photoactivation (**Supplementary Fig. 6e**)^14^. Glucose addition to glucose-deprived cells stimulated glycolysis and lactate production^38^, increasing fluorescence in both series (**Supplementary Fig. 6f,g**).

**Supplementary Table 1.**
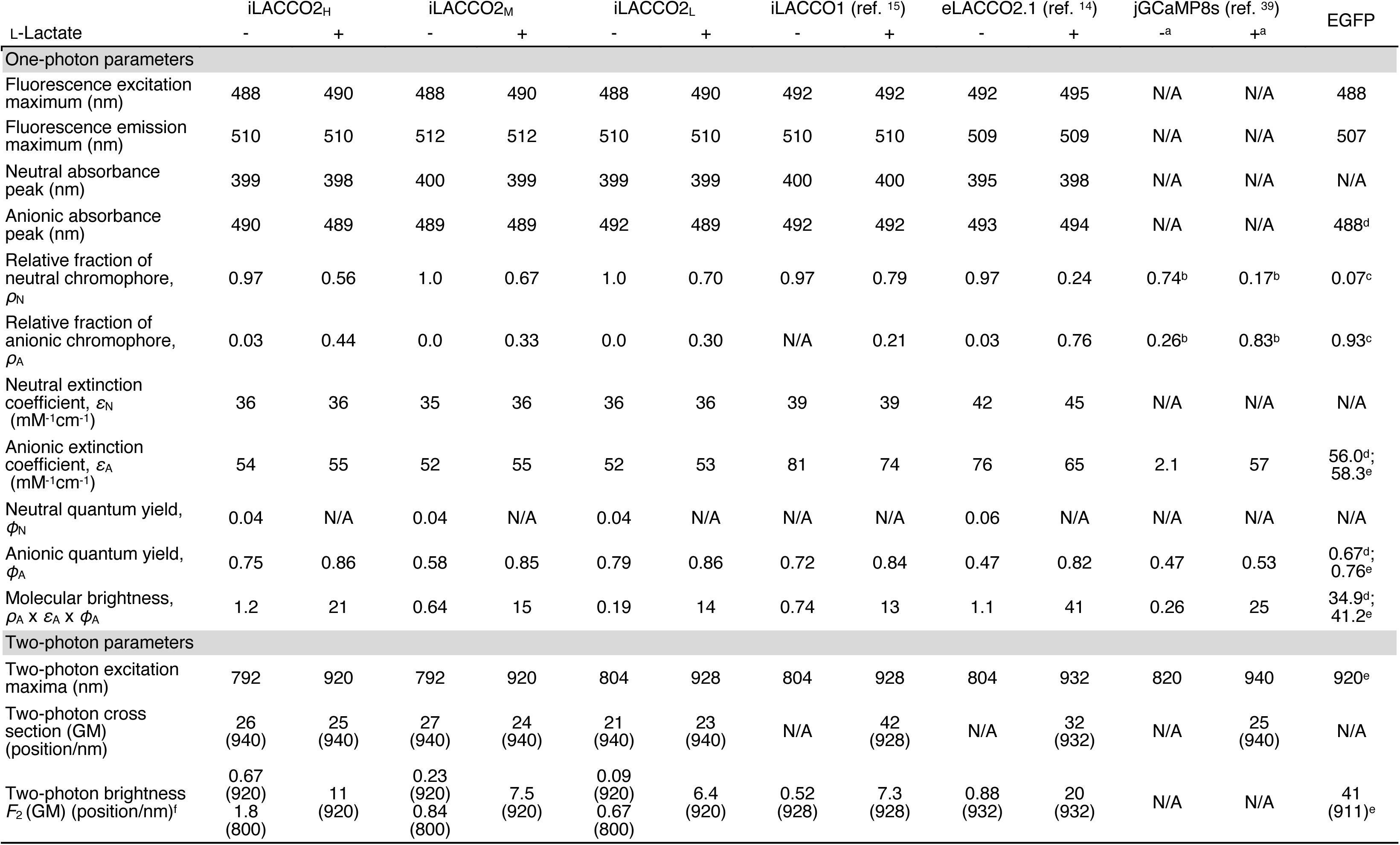
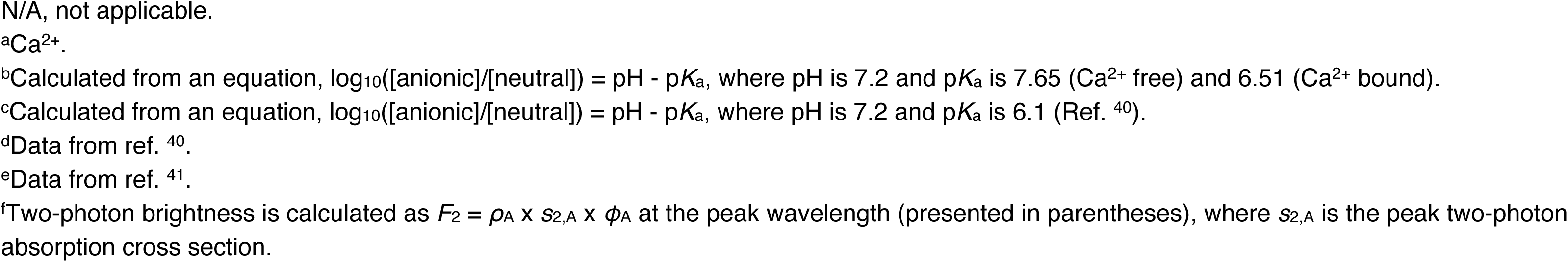
One- and two-photon photophysical parameters of iLACCO2 variants.

**Supplementary Table 2.**
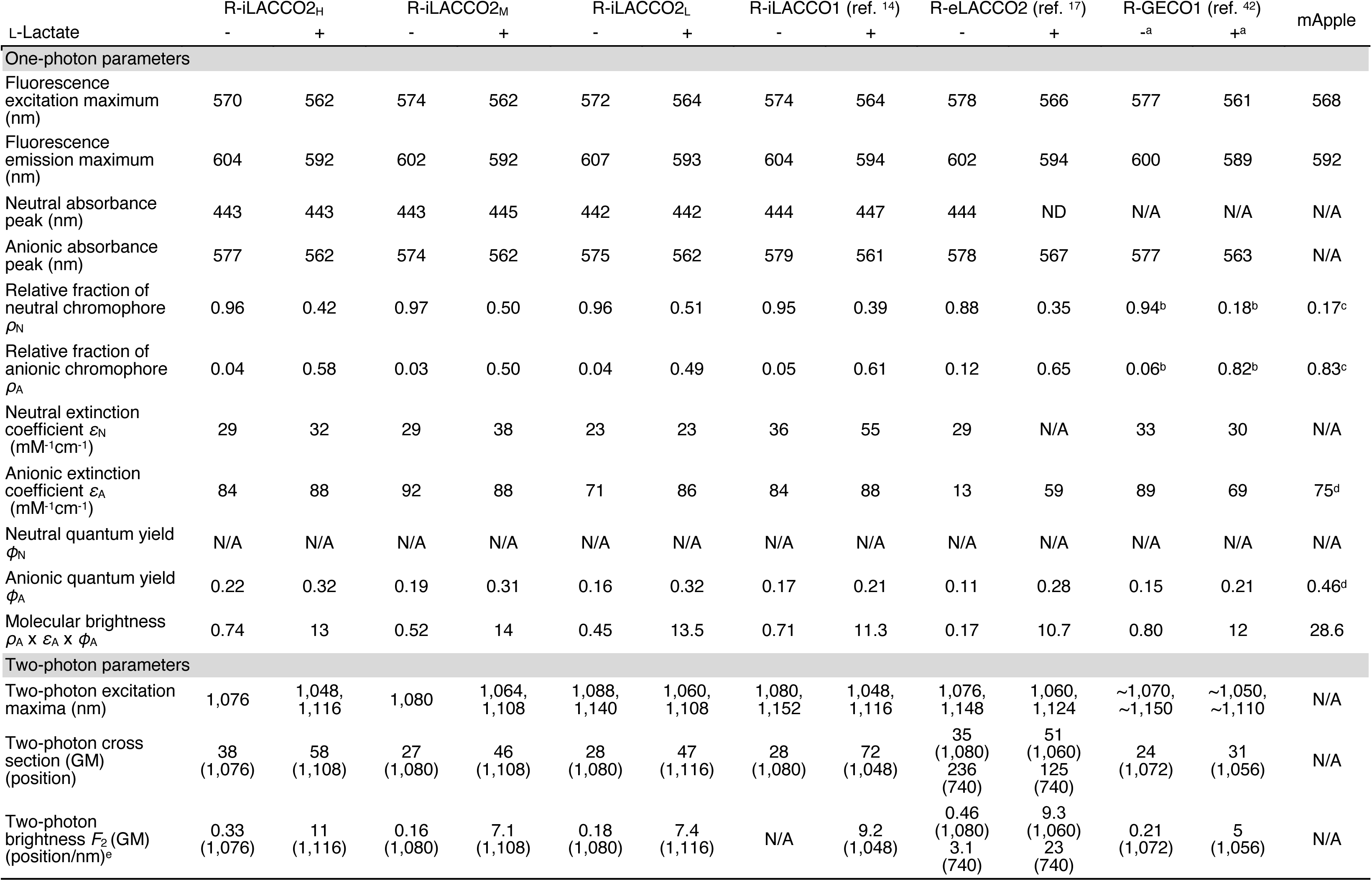
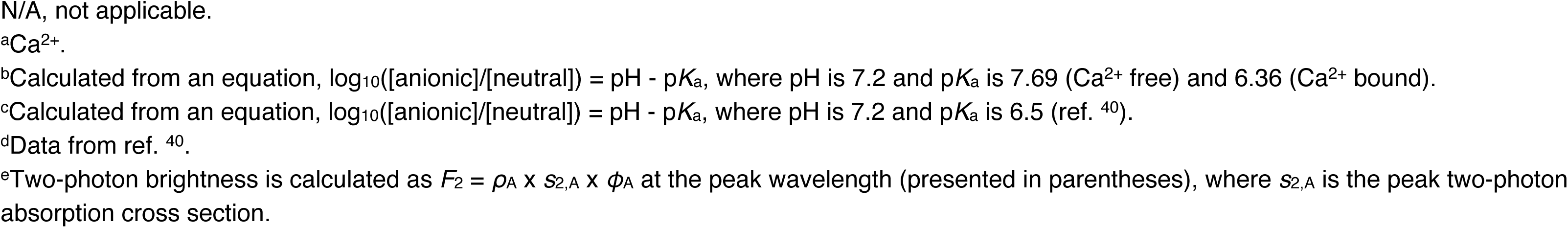
One- and two-photon photophysical parameters of R-iLACCO2 variants.

